# Conserved copper(I)-binding auxiliary domains link virulence-associated LPMOs to copper homeostasis in the intestinal environment

**DOI:** 10.64898/2026.08.05.742968

**Authors:** Eirik G. Kommedal, Hanne Berggreen, Synnøve Elisa Rønnekleiv, Henrik Vinther Sørensen, Ane Runningen, Yong Zhou, Ute Krengel, Åsmund K. Røhr, Vincent G. H. Eijsink, Zarah Forsberg

## Abstract

Multidomain lytic polysaccharide monooxygenases (LPMOs) are widely distributed in human pathogenic bacteria and increasingly recognized for promoting host interactions and virulence. Recent work has shown that immunizing mice with CbpD, a trimodular LPMO from *Pseudomonas aeruginosa*, provides protection against lethal *P. aeruginosa* infection. In several human intestinal pathogenic bacteria, LPMOs occur as tetra- and pentamodular proteins and their ability to interact with the host and facilitate virulence has primarily been ascribed to their ability to bind to *N*-acetylglucosamine containing glycans. Recently, AlphaFold predictions and substrate-binding studies of the *Vibrio cholerae* colonization factor GbpA suggested the presence of a copper-binding site in its non-catalytic third domain. In light of the recent demonstration that MUC2, the primary intestinal mucin, harbors two conserved and distinct copper binding sites for Cu(II) and Cu(I), we hypothesized that the auxiliary non-catalytic copper-binding domains of intestinal LPMOs evolved as a response to the intestinal environment. Here, we examine GbpA and homologous multidomain LPMOs from food-borne intestinal disease-causing Gram-positive bacteria from the genera *Bacillus* and *Listeria*. By combining mutagenesis with biochemical, spectroscopic, and computational approaches, we demonstrate that these multidomain LPMOs contain conserved, yet structurally distinct copper(I)-binding motifs on their non-catalytic third domain. A comprehensive review and reassessment of the literature spanning the past two decades reveals consistent links between the copper-binding ability of these LPMOs and host-pathogen interactions. Thus, our findings link these LPMOs to copper management in the intestine, offering an explanation for why their unique architecture is preserved across different bacterial lineages causing intestinal disease.

## Introduction

Pathogenic species within the *Vibrio*, *Bacillus*, and *Listeria* genera share the ability to colonize the intestinal tract and cause severe disease in humans and animals. The aquatic Gram-negative bacterium *Vibrio cholerae* is a food-borne pathogen and the etiologic agent of cholera, a waterborne diarrheal disease responsible for an estimated ∼ 4 million cases and up to 143,000 deaths globally each year [1], whilst the foodborne Gram-positive pathogens *Bacillus cereus* and *Listeria monocytogenes* cause gastrointestinal illness [2] and listeriosis [3], respectively. Several chitin-degrading enzymes, including chitin-active lytic polysaccharide monooxygenases (LPMOs), have been implicated in host-pathogen interactions [4–10], suggesting that these enzymes may play dual roles, in nutrient acquisition and in host colonization and virulence. During infection, these pathogens will encounter the intestinal mucus layer, which does not contain chitin, but does contain glycan structures that include the building block of chitin, *N*-acetylglucosamine.

The intestinal mucus layer, primarily composed of the hydrogel forming MUC2 mucin, separates the luminal space from the epithelial cells and serves as the primary interaction site for pathogenic bacteria [11–13]. The N-terminal D1 region of MUC2 is a copper chaperone that contains two conserved copper-binding sites coordinating Cu(II) and Cu(I), respectively. Thus, MUC2 plays a role in copper management in the intestine, likely preventing copper toxicity, while facilitating uptake of this important trace metal into cells [14–16]. How exactly this copper-binding capacity of MUC2 affects pathogenic intestinal bacteria remains unknown. It is worth noting though that it has recently been shown that copper promotes the activity of alpha-kinase 1 (ALPK1), a cytosolic master regulator of host cell defense against bacterial infection, and that copper promotes innate immunity [17]. Interestingly, a well-known virulence factor of *Vibrio cholerae* known as “colonization factor” or *N*-acetylglucosamine binding protein A (GbpA or *Vc*GbpA), is a copper-binding protein. This four-domain protein has a catalytically active, copper-binding N-terminal LPMO domain [18] and is known to facilitate attachment of bacterial cells to intestinal cells [8, 9] and to stimulate mucin production [19]. Recent data suggest that the third domain of *Vc*GbpA has an additional copper site [20, 21]. Strikinglye, other intestinal pathogens, including Gram-positive *Bacillus cereus* and *Listeria monocytogenes*, have LPMOs with similar modular architectures (*vide infra*).

LPMOs are copper-dependent enzymes that catalyze oxidative cleavage of insoluble polysaccharides, thus generating amorphicity and new access points for hydrolytic enzymes [22, 23]. These secreted enzymes coordinate a single copper ion through two strictly conserved histidine residues, including an N-terminal histidine that emerges upon cleavage of the signal peptide [24, 25]. Their catalytic cycle involves reduction of the active site copper from Cu(II) to Cu(I), enabling a reaction with H_2_O_2_ to form a highly reactive copper-oxyl species that abstracts a hydrogen atom from the polysaccharide substrate [26–32]. Like other redox-active metalloenzymes, LPMOs are susceptible to oxidative damage; in the absence of a suitable substrate, reactive intermediates can engage in off-pathway reactions that damage residues in and around the active site leading to irreversible enzyme inactivation and loss of the copper cofactor [26, 33, 34]. Consequently, LPMO function and stability depend on copper availability, redox conditions, and substrate interactions, all of which might be limiting in host environments.

LPMOs occur as single-domain or multidomain proteins. Multidomain architectures that pair an LPMO with a carbohydrate-binding module (CBM) and/or a glycoside hydrolase (GH) of matching specificity likely reflect roles in biomass degradation [35–37]. On the other hand, the biological role of a distinct set of multidomain LPMOs commonly found in pathogenic bacteria (Figure 1) is less clear. These LPMOs typically contain one or two internal domains of unknown function positioned between an N-terminal chitin-active LPMO domain and one or two C-terminal CBMs (typically chitin-binding modules of families CBM5/12 or CBM73), and are frequently annotated as GlcNAc-binding protein A (GbpA) [8] or chitin-binding protein (CBP) [38–40]. Their ecological distribution and expression patterns appear uncoupled from the presence of chitin and seem associated with conditions encountered during host colonization and infection [5, 7–9, 19], raising the question of whether chitin is a biologically relevant substrate for these proteins.

**Figure 1.**
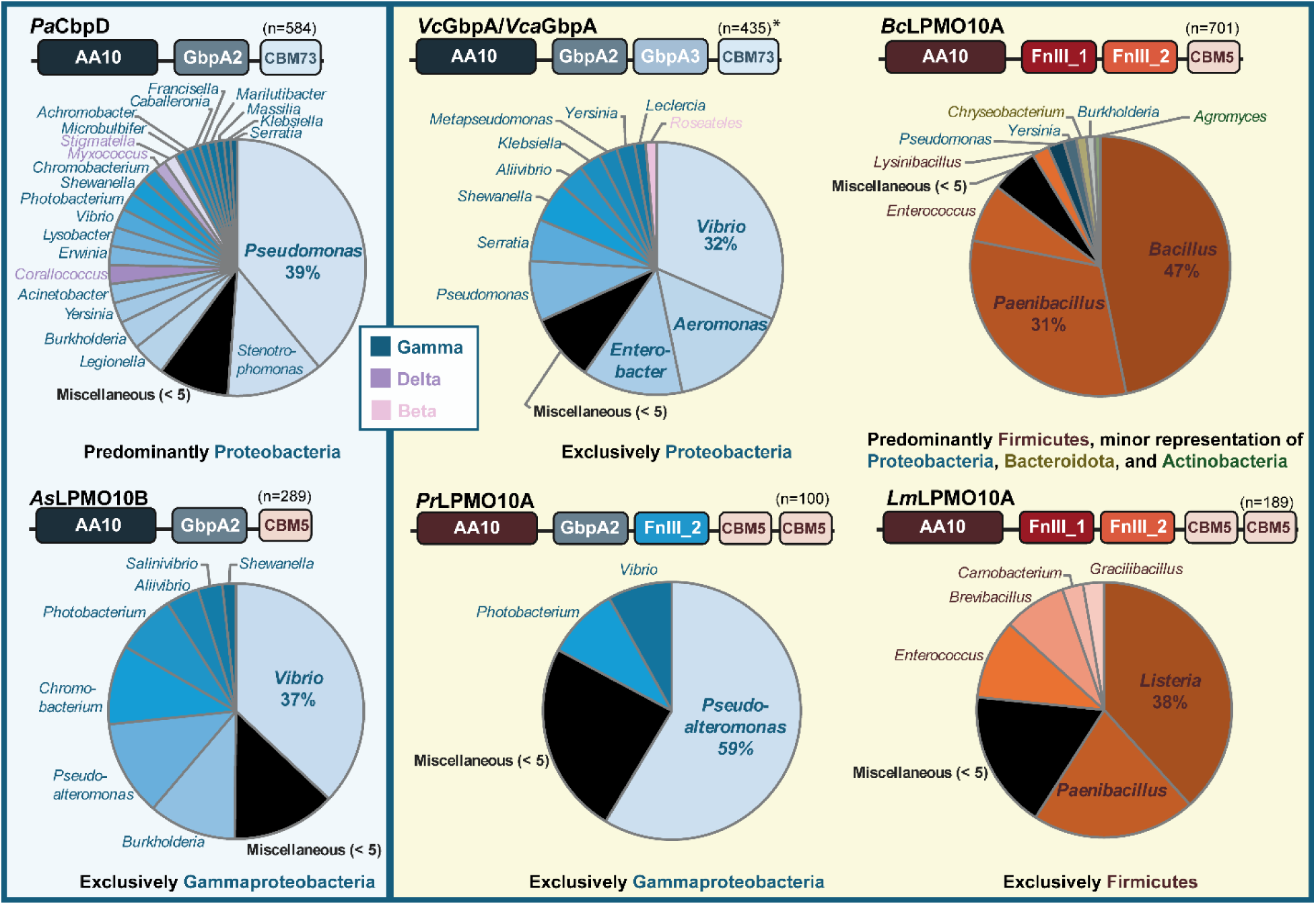
Taxonomy and domain architectures of prevalent multimodular LPMOs. The six multimodular LPMO architectures shown here all rank among the ten most common architectures identified in InterPro (IPR004302; 197 architectures in total, December 2025). Gram-negative bacteria are indicated in blue and Gram-positive bacteria in rust. Architectures on the left contain a single internal auxiliary domain (GbpA2), whereas those on the right contain two auxiliary domains (named GbpA2 and GbpA3 or FnIII-like domains 1 and 2). The domain structures of representative proteins are shown above each group: *Pa*CbpD (*Pseudomonas aeruginosa*) [5], *As*LPMO10B (*Aliivibrio salmonicida*) [6], *Vc*GbpA/*Vca*GbpA (*Vibrio cholerae/Vibrio campbellii*) [42, 43], *Pr*LPMO10A (*Pseudoalteromonas rubra*; uncharacterized), *Bc*LPMO10A (*Bacillus cereus*) [44], and *Lm*LPMO10A (*Listeria monocytogenes*) [45]. Sequences were grouped by phylum or class when possible; CbpD-like and GbpA-like proteins span Gamma-, Delta-, and Betaproteobacteria, although Gammaproteobacteria predominate. An asterisk above the number of GbpA-like sequences indicates that 11 of 435 sequences contain a CBM5 instead of the CBM73. The “Miscellaneous” category groups genera represented by fewer than five sequences in the dataset.

Despite being widespread in pathogenic bacteria, and despite functional information that links *Vc*GbpA and a trimodular LPMO from the airway pathogen *P. aeruginosa* [41] to pathogenicity, the biological roles of these multidomain LPMOs, including the role of their auxiliary internal domains (Figure 1), remain poorly understood. Recent work on mucins [14–16] and on the regulation of innate immunity [17], suggest that copper and mucosal copper-binding sites may be important determinants of intestinal infections. Furthermore, recent work on multidomain LPMOs from two Gram-negative pathogens suggests that these enzymes may contain copper binding sites in their auxiliary domains [20, 21], with a role in regulating LPMO activity and/or sequestering copper. Using phylogenetic analysis as well as site-directed mutagenesis and functional studies of three different multidomain LPMOs from the intestinal bacterial pathogens *Vibrio cholera*, *Bacillus cereus*, and *Listeria monocytogenes*, we show that one of the internal domains of these proteins, the third domain, harbors a copper(I)-binding site. These copper binding domains have a fibronectin-like fold and while the various LPMOs carry distinct copper-binding motifs in this domain, such motifs are only found in fibronectin domains attached to LPMOs. Thus, our results show that the ability to sequester copper(I) is a conserved trait in the multidomain LPMOs of Gram-negative and Gram-positive intestinal pathogens (but not in airway pathogens; *vida infra*). Based on these results and a critical reassessment of literature data, we propose that the role of these unique LPMOs in intestinal disease relates to their impact on copper management in the intestine.

## Results and discussion

### Discovery of a conserved metal binding function in FnIII-like domains exclusively tethered to chitin-oxidizing LPMOs from pathogenic bacteria

In our recent comparative study of GbpA proteins from *V. cholerae* and *V. campbellii*, we identified a conserved set of residues on the GbpA3 domain that putatively interact with the Cu-containing active site and the substrate binding surface of the LPMO domain under reducing conditions [21]. Following up on this work, we revisited previous work on multidomain LPMOs from pathogenic Gram-positive bacteria: the four-domain LPMO from *B. cereus*, *Bc*LPMO10A [44], and the five-domain LPMO from *Listeria monocytogenes*, *Lm*LPMO10A [45] (Figure 1). In addition, we carried out site-directed mutagenesis studies to experimentally assess predicted copper-binding abilities.

Structure prediction of *Bc*LPMO10A and *Lm*LPMO10A with AlphaFold3 revealed that, also in these proteins, the third domain may interact with the active site copper in the catalytic domain (Figure 2). In the case of *Lm*LPMO10A, copper alone was sufficient to trigger the predicted conformational change that promotes interaction between the LPMO domain and the third domain. In contrast, for *Bc*LPMO10A, such a conformational change was observed only when, next to a copper ion, two calcium ions, one coordinated in each internal FnIII-like domain, were included in the AlphaFold3 structure prediction (Figure S1). This observation led us to identify a conserved putative calcium-binding site in both FnIII-like domains in *Bc*LPMO10A-like (Figure S2) and *Lm*LPMO10A-like (Figure S3) enzymes (Figure 2). Notably, these calcium-binding motifs are less well conserved in FnIII-like domains associated with modular GH18 chitinases (Figure S4). It is also worth noting that, while clearly conserved in *Bc*LPMO10A-like and *Lm*LPMO10A-like enzymes, calcium binding is not predicted for the GbpA2 and GbpA3 domains of *Vc*GbpA and *Vca*GbpA; accordingly, addition of calcium ions did not affect the outcome of the AlphaFold3 predictions for these proteins.

**Figure 2.**
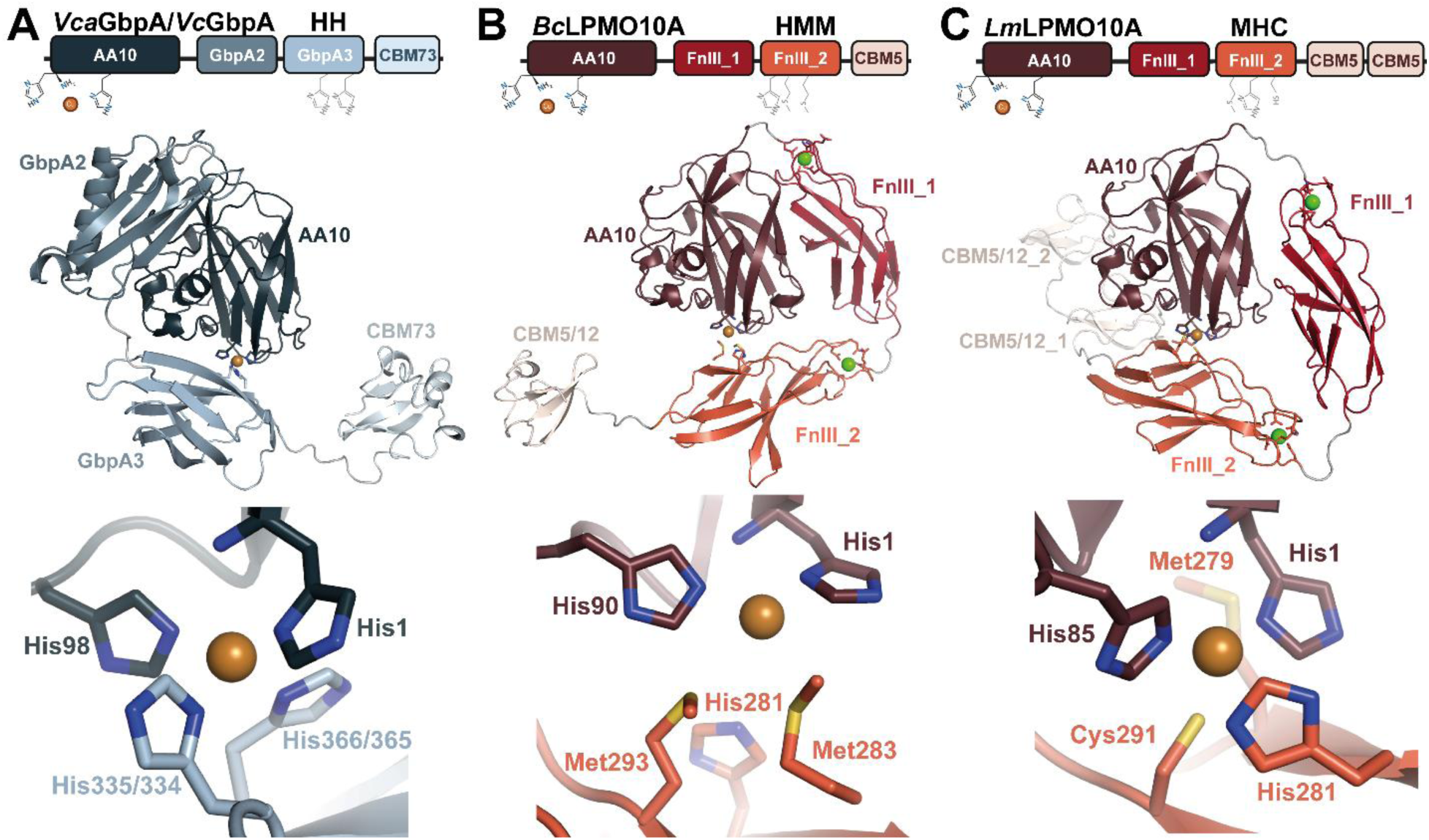
Domain architecture and predicted structures of *Vca*GbpA, *Bc*LPMO10A, and *Lm*LPMO10A. Panel A shows *Vca*GbpA/*Vc*GbpA, panel B shows *Bc*LPMO10A, and panel C shows *Lm*LPMO10A. In each panel, the top part shows the domain organization, highlighting the His-brace copper site in the catalytic AA10 domain and the predicted copper-binding residues in the third domain. The middle part illustrates predicted interdomain interactions between the domains. Copper ions are shown as orange spheres. Two Ca^2+^ were included in the predictions for *Bc*LPMO10A and *Lm*LPMO10A, but not for *Vca*GbpA/*Vc*GbpA; calcium ions are shown as green spheres. The bottom part provides a close-up view of the residues coordinating the copper in the “closed” structure. While these residues differ between the three enzymes, they are largely conserved in LPMOs with similar domain architectures (see Figures S2 and S3 for the FnIII-like domains of *Bc*LPMO10A and *Lm*LPMO10A and Zhou et al. [21] for GbpA3). Panel A shows *Vca*GbpA but is representative of *Vc*GbpA due to high (82%) sequence similarity. Residue numbering in the His-brace is identical, whereas both numberings are shown for histidines in the GbpA3 domain. All structures were predicted using AlphaFold3 [46].

FnIII domains display low sequence identity (∼30% or lower) but share a highly conserved structural framework. These domains have been described as mediators of protein-protein interactions and as “spacers”, positioning functional modules in the appropriate spatial context needed to achieve their biological roles [47, 48]. Notably, the GbpA3 domain has a similar fold and topology (Figure S5), suggesting that GbpA3 actually is an FnIII-like domain. Still, the domains differ, since the GbpA3 domain does not seem to bind calcium and since the copper binding residues are located very differently (*vide infra*). Interestingly, like GbpA3 from *Vc*GbpA and *Vca*GbpA [21], the FnIII_2 domains in *Bc*LPMO10A and *Lm*LPMO10A contain conserved copper-binding motifs comprised of His281–Met283–Met293 and Met279–His281–Cys291, respectively (Figure 2 and Figures S2 & S3). These putative copper sites in the FnIII_2 domains share a common primary amino acid sequence starting close to the N-terminus: H^321^X^322^M^323^ … M^333^W^334^ and M^279^X^280^H^281^ … C^291^W^292^, for *Bc*LPMO10A and *Lm*LPMO10A, respectively. The tryptophan residue is conserved in both the FnIII_1 and FnIII_2 domains of *Bc*LPMO10A-like (Figure S2) and *Lm*LPMO10A-like proteins (Figure S3), as well as in the FnIII domains of modular GH18s (Figure S4), but the putative copper-binding residues only appear in the FnIII_2 like domains of the LPMOs. This suggests that the copper binding motif in these FnIII-like domains co-evolved with the LPMO domain. Interestingly, a subpopulation of the *Bc*LPMO10A-like LPMOs, all from *Paenibacillus larvae* and including *Pl*CBP49 [40] contain an RMM motif (Figure S6). To our knowledge, an RMM motif has never been reported to bind copper as confirmed by experimental data shown below. It is noteworthy that this four-domain LPMO is a key virulence factor in American foulbrood of honeybees and thus occurs in a chitin-rich environment [40].

### Probing for an interaction between the LPMO domain and the third domain in *Vca*GbpA, *Bc*LPMO10A and *Lm*LPMO10A

Earlier studies of the putative copper binding abilities of the auxiliary GbpA3 domain in *Vc*GbpA/*Vca*GbpA [21] and a GbpA-like protein from *E. cloacae* [20], have focused on the possible interaction between this domain and the catalytic domain suggested by the AlphaFold3 predictions. It has been speculated that such an interaction could provide a regulatory mechanism: in the absence of substrate the GbpA3 domain interacts with the reduced LPMO domain, preventing the LPMO from engaging in off-pathway reactions that would result in enzyme damage; in the presence of substrate, the CBM binds to the substrate, which changes interdomain interactions and enables the LPMO active site to engage in productive substrate oxidation [20, 21]. So far, the most convincing experimental proof for this interaction is based on studies of binding of reduced truncated variants of *Vc*GbpA and *Vca*GbpA to β-chitin [21]. These studies showed that the CBM-truncated variants bind much less to chitin than the full-length enzyme and the isolated LPMO catalytic domain, which is compatible with the idea that the LPMO substrate-binding surface is blocked by the interaction with the GbpA3 domain in the CBM-truncated variant.

To assess this interaction, we generated variants of the CBM-truncated constructs of *Vca*GbpA, *Bc*LPMO10A and *Lm*LPMO10A in which the copper binding residues in the third domain were replaced by alanine, assuming that these substitutions would weaken the domain-domain interaction (Table 1). Instead of *Vc*GbpA, we used highly similar *Vca*GbpA because we obtained better expression for the CBM-truncated variant of this enzyme. Of note, CBM-truncated variants were used as the CBMs bind strongly to chitin and we would not expect to see any differences between wildtype and alanine variants. Although, subsequent binding studies with β-chitin under reducing and non-reducing conditions showed variability between the enzymes, clear trends were visible: the alanine variants bind better to chitin than the wildtype enzymes and this effect is more pronounced under reducing conditions (Figure 3). This observation provides experimental proof for the hypothesis that the two domains interact and that the copper binding ability of the third domain is a mediator of this interaction.

**Figure 3.**
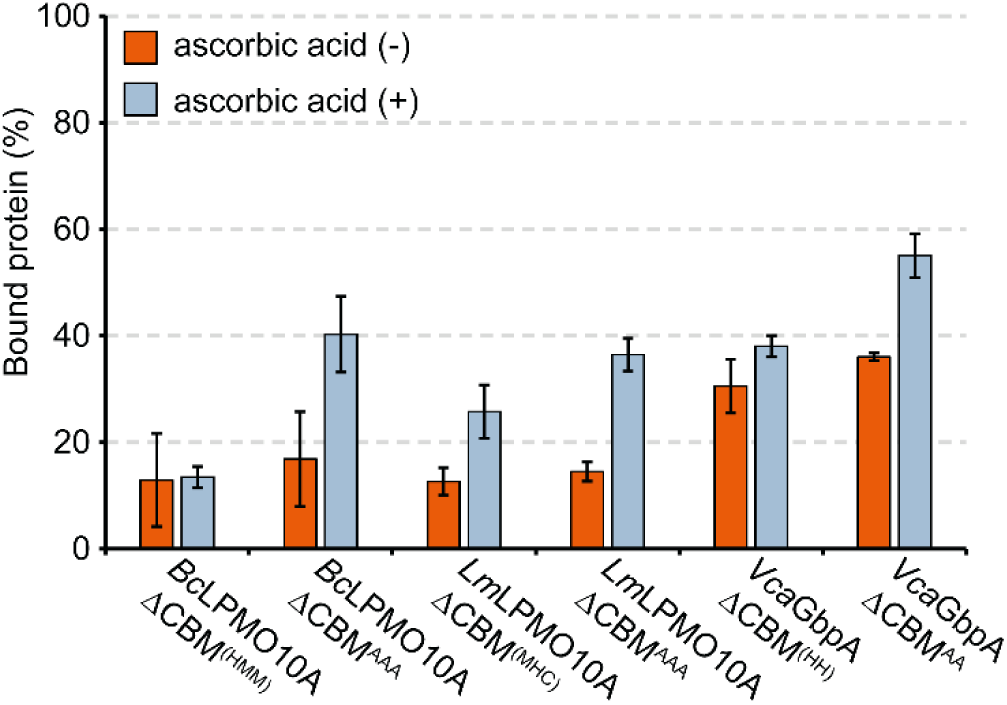
Binding of CBM-truncated variants of *Vca*GbpA, *Bc*LPMO10A and *Lm*LPMO10A to chitin. Cu(II)-saturated *Vca*GbpA-ΔCBM^(HH)^, *Vca*GbpA-ΔCBM^AA^, *Bc*LPMO10A-ΔCBM^(HMM)^, *Bc*LPMO10A-ΔCBM^AAA^, *Lm*LPMO10A-ΔCBM^(MHC)^ and *Lm*LPMO10A-ΔCBM^AAA^ were incubated with 10 g/L β-chitin in the absence (orange) and presence (light blue) of 1 mM ascorbic acid in 50 mM Tris (pH 7.5) at 22 °C, 800 rpm for 30 min. Subsequently, the chitin-bound enzyme fraction was separated from the unbound fraction by filtration using a vacuum manifold. The non-bound enzyme fraction was quantified using a standard Bradford assay. Error bars represent standard deviations (n = 3).

**Table 1.** Overview of enzyme variants used in this study. The proteins studied were *Bc*LPMO10A (UniProt ID: Q81CG6), *Lm*LPMO10A (UniProt ID: Q8Y4H4), *Vca*GbpA (UniProt ID: A7N3J0), and *Vc*GpbA (UniProt ID: Q9KLD5). Full-length proteins are denoted FL and CBM truncated variants are denoted ΔCBM. Amino acid numbering relates to the sequence of the native protein excluding the native signal peptide, *i.e.,* the first histidine is His1. Wildtype sequence motifs related to copper binding by the third domain appear between parentheses, whereas mutated motifs appear without parentheses.

| Enzyme variant | Substitution(s) | Residues |
| --- | --- | --- |
| ***Bacillus cereus*** |  |  |
| FL<sup>(HMM)</sup> | - | 1-415 |
| FL<sup>AAA</sup> | H281A, M283A, M293A | 1-415 |
| ΔCBM<sup>(HMM)</sup> | - | 1-364 |
| ΔCBM<sup>AAA</sup> | H281A, M283A, M293A | 1-364 |
| FnIII_2<sup>(HMM)</sup> | - | 270-364 |
| FnIII_2<sup>AAA</sup> | H281A, M283A, M293A | 270-364 |
| FnIII_2<sup>RMM</sup> | H281R | 270-364 |
| ***Listeria monocytogenes*** |  |  |
| FL<sup>(MHC)</sup> | - | 1-451 |
| FL<sup>AAA</sup> | M279A, H281A, C291A | 1-451 |
| ΔCBM<sup>(MHC)</sup> | - | 1-355 |
| ΔCBM<sup>AAA</sup> | M279A, H281A, C291A | 1-355 |
| FnIII_2<sup>(MHC)</sup> <sup>a</sup> | - | 268-355 |
| FnIII_2<sup>AAA</sup> <sup>a</sup> | M279A, H281A, C291A | 268-355 |
| ***Vibrio campbellii*** |  |  |
| FL<sup>(HH)</sup> | - | 1-464 |
| FL<sup>AA</sup> | H335A, H366A | 1-464 |
| ΔCBM<sup>(HH)</sup> | - | 1-392 |
| ΔCBM<sup>AA</sup> | H335A, H366A | 1-392 |
| GbpA3<sup>(HH)</sup> <sup>a</sup> | - | 295-392 |
| GbpA3<sup>AA</sup> <sup>a</sup> | H335A, H366A | 295-392 |
| ***Vibrio cholerae*** |  |  |
| GbpA3<sup>(HH)</sup> | - | 294-391 |
| GbpA3<sup>AA</sup> | H334A, H365A | 294-391 |

<sup>a</sup> Successful production of these proteins was not achieved.

Possible domain interactions were also assessed by using small-angle X-ray scattering (SAXS) to study the solution structures of the copper-saturated full-length wildtype proteins, i.e., *Vca*GbpA^(HH)^, *Bc*LPMO10A^(HMM)^, and *Lm*LPMO10A^(MHC)^, as well as non-copper saturated (*apo*) and copper-saturated (*holo*) CBM-truncated *Bc*LPMO10A^(HMM)^ under aerobic conditions in the absence or presence of a reductant. Under all conditions assessed here, all four proteins exhibited extended conformations, as has previously been shown for the *apo*-form and Cu(II)-state of *Vc*GbpA [42, 49] and *Vca*GbpA [43] (Figure S7). Thus, we were not able to prove an interaction between the LPMO domain and the GbpA3/FnIII_2 domain with SAXS.

### FnIII-like domains with metal binding motifs bind copper with a preference for Cu(I)

AlphaFold3 was used to investigate potential copper-binding sites in the isolated third domains of *Bc*LPMO10A, *Lm*LPMO10A, and *Vc*GbpA/*Vca*GbpA, thereby avoiding interference from the catalytic LPMO copper site. All domains were predicted to coordinate copper in these isolated domain models. To further evaluate these sites, geometry optimization was performed using the extended semi-empirical tight-binding method (GFN2-xTB) [50] with the ALPB implicit water solvent model. Calculations were performed for the isolated FnIII-like domains (residues 294–391 in *Vc*GbpA, 270-364 in *Bc*LPMO10A, and 268–355 in *Lm*LPMO10A) including Cu(I). The optimized structures of *Bc*FnIII_2 and *Lm*FnIII_2 display distorted trigonal planar Cu(I) geometries, with Cu– N/S bond lengths of 2.0–2.4 Å (Figure 4A), consistent with reported Cu(I)-binding sites [51–53] (see Figure S8 for detailed geometries and comparisons). In contrast, the two histidines predicted to coordinate copper in *Vc*GbpA3 form a bent two-coordinate copper site, deviating from the linear geometry typically preferred for two-coordinate Cu(I) centers [54]. While linear coordination is readily achieved in small-molecule Cu(I) complexes, distorted geometries are not uncommon in proteins, where constraints imposed by the protein scaffold and the fixed positioning of coordinating residues can prevent the formation of an ideal linear Cu(I) center [55].

**Figure 4.**
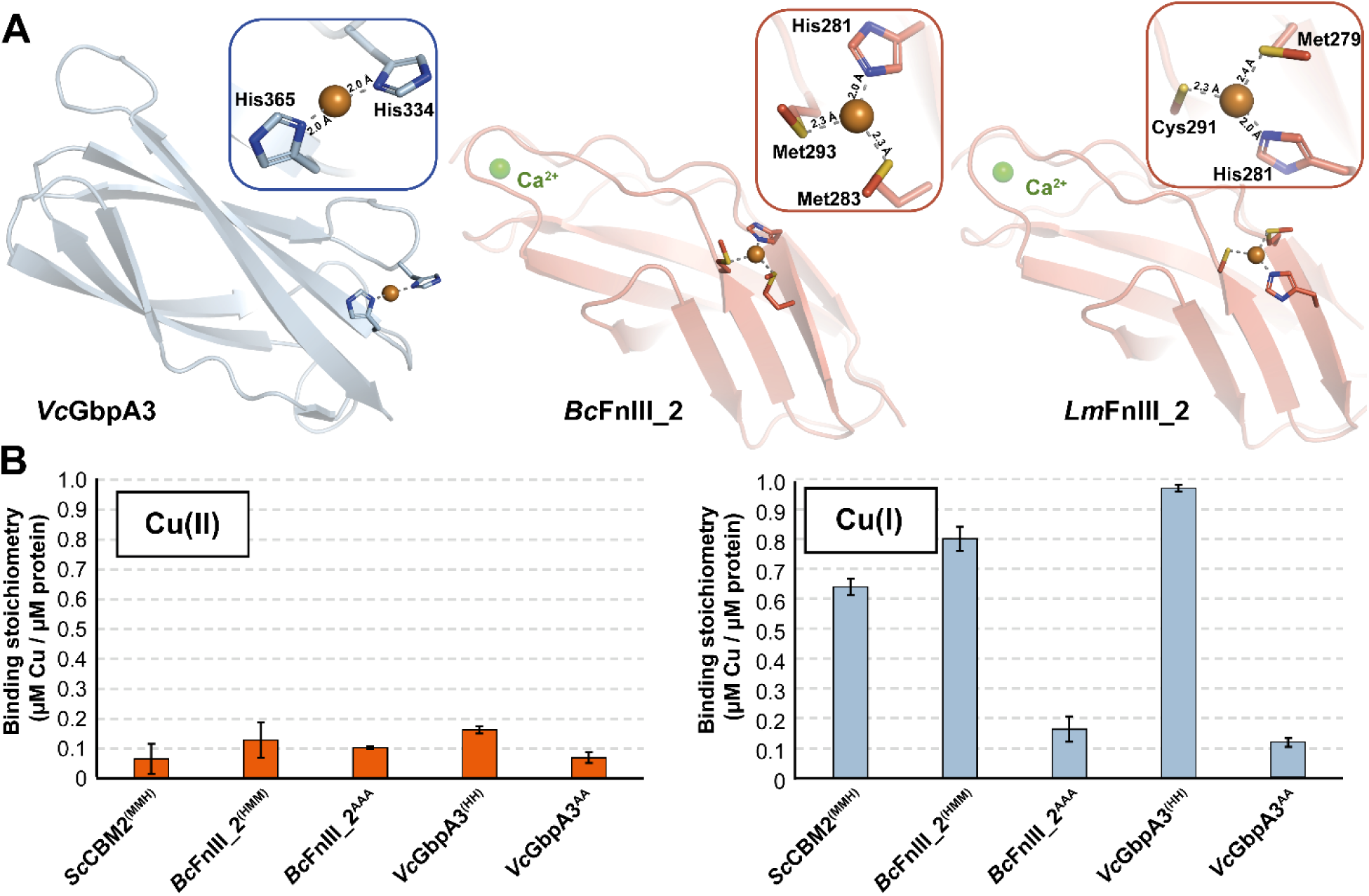
Copper binding by wildtype and alanine-substituted variants of the *Bc*FnIII_2 and *Vc*GbpA3 domains. Panel A shows geometry optimized models of the copper binding sites in *Vc*GbpA3^(HH)^, *Bc*FnIII_2^(HMM)^ and *Lm*FnIII_2^(MHC)^. Figure S8 provides additional details. Panel B shows the analysis of copper binding by wildtype and alanine-substituted variants of *Bc*FnIII_2 and *Vc*GbpA3 under oxidizing (left) and reducing (right) conditions. *Sc*CBM2, containing a Met–Met–His Cu(I)-binding motif [52], was used as a positive control for Cu(I)-binding. Each protein (4 µM) was incubated with 8 µM CuSO4 for 10 min in sodium phosphate buffer (pH 6.0), either with or without 20 µM ascorbic acid. Protein was removed by ultrafiltration (3 kDa MWCO) and the filtrate, containing unbound copper, was mixed with bathocuproine disulfonate (BCS) and 40 µM ascorbic acid to ensure complete reduction and formation of the Cu(I)–BCS complex. Fluorescence (Ex/Em = 290/325 nm) was recorded and converted to copper concentration using a CuSO4 standard curve (0–8 µM). The y-axis represents the amount of bound Cu(II) or Cu(I), normalized to 1 µM of protein, indicating the apparent binding stoichiometry for each condition. Error bars represent standard deviations (n = 3).

To investigate the copper binding ability of the two GbpA3 domains (*Vc* & *Vca*) and the two FnIII_2 domains (*Bc* & *Lm*), these domains were produced as single domains, which, despite extensive efforts, was successful in only two cases, *Vc*GbpA3 (with a HH motif) and *Bc*FnIII_2 (with a HMM motif). For these two single domains, we also created variants in which residues predicted to be involved in copper coordination were mutated to alanine (Table 1). In addition, the HMM motif in *Bc*FnIII_2 was mutated to the RMM motif found in *P. larvae* CBP49. Binding of Cu(I) and Cu(II) by these domains was assessed using bathocuproine disulfonate (BCS) as a probe [56, 57], whereas binding of Cu(II) was also assessed using electron paramagnetic resonance (EPR) spectroscopy. For the BCS assays, the CBM2 domain of LPMO10C from *Streptomyces coelicolor* was used as positive control for Cu(I) binding as this CBM was recently shown to bind Cu(I) through a MMH motif [52].

The results show that *Bc*FnIII_2^(HMM)^ and *Vc*GbpA3^(HH)^ bind reduced copper (Cu(I)), whereas alanine substitution of the coordinating residues abolishes copper binding (Figure 4B). In contrast, Cu(II) binding was negligible for all tested variants. The absence of Cu(II) binding by FnIII_2 and GbpA3 is further supported by EPR measurements (Figure S9) of full-length enzymes (*Vca*GbpA^(HH)^, *Vca*GbpA^AA^, *Bc*LPMO10A^(HMM)^, *Bc*LPMO10A^AAA^, *Lm*LPMO10A^(MHC)^ and *Lm*LPMO10A^AAA^), as the spectra of wildtypes and alanine substituted variants, i.e., variants lacking the copper-binding site in the third domain, were indistinguishable. Binding of Cu(II) to the third domain in the wildtype enzymes would have yielded an additional Cu(II) signal that should not be present in the alanine substituted variants.

AlphaFold3 models did not support copper binding at the RMM site in *Pl*CBP49, neither in the isolated FnIII_2-like domain nor in the full-length enzyme, and no interdomain interaction was predicted. Consistently, AlphaFold did no longer predict an interdomain interaction after introduction of the RMM motif into *Bc*LPMO10A. To experimentally assess the contribution of the RMM motif, we mutated *Bc*FnIII_2 (HMM→RMM) and examined copper binding using the BCS assay (Figure S10). This substitution resulted in a loss of copper binding, comparable to the loss seen in the *Bc*FnIII_2^AAA^ variant, indicating that the histidine residue is a primary contributor to the copper affinity of this site and showing that the third domain of *Pl*CBP49, which is not from an intestinal bacterial pathogen, likely does not bind copper.

### Copper binding FnIII-like domains prevent ascorbic acid oxidation and limit off-pathway copper reactivity

To assess how these copper binding domains influence ascorbic acid oxidation in the presence and absence of an LPMO substrate, β-chitin, we employed a luminescent terbium–2,6-dipicolinic acid (TbDPA) assay. This approach exploits the ability of ascorbic acid to quench TbDPA luminescence, enabling indirect quantification of ascorbic acid consumption [58]. As ascorbic acid is oxidized, quenching decreases and luminescence intensity increases accordingly. Importantly, this method overcomes the strong optical interference caused by chitin particles at 255 nm and allows monitoring of the consumption of ascorbic acid directly in substrate-containing reactions.

In the absence of a carbohydrate substrate, LPMO-mediated oxidation of ascorbic acid and concomitant formation of H_2_O_2_ promote the off-pathway peroxidase reaction that may damage the active-site histidine brace, resulting in copper loss and enzyme inactivation [26, 33, 59]. Enzyme inactivation and ascorbic acid depletion are self-reinforcing processes: as the enzyme inactivates, copper is released from its catalytic center, which leads to faster (copper-catalyzed) oxidation of ascorbic acid and formation of H_2_O_2_, which again leads to faster enzyme inactivation [60]. These effects are prevented by the presence of substrate, when the oxidase reaction is less prominent and the resulting H_2_O_2_ is used productively, and/or if copper released from damaged active sites is sequestered, for example by the GbpA3 or FnIII_2 domains of the multimodular LPMOs studied here.

The results, obtained with the full-length enzymes, show more rapid depletion of ascorbic acid for the alanine-substituted variants compared to their wildtype counterparts regardless of the presence of β-chitin, and also show that β-chitin, expectedly, dampens ascorbic acid consumption for all enzymes (Figure 5). These effects are most prominent for *Vca*GbpA and *Bc*LPMO10A, and less so for *Lm*LPMO10A. Comparison of the wildtype enzymes (Figure 5A,C,E) shows that the enzymes have different life times in the absence of substrate, which is a consequence of different abilities to (a) protect the catalytic domain from engaging in off-pathway reactions through domain-domain interactions, (b) different abilities to prevent damage through hole hopping mechanisms [61, 62] and/or (c) different abilities to sequester free Cu(I). This comparison also shows that the impact of the substrate, present at a non-saturating low concentration of 2 g/L, differs between the enzymes, which suggests different abilities to interact with and oxidize chitin, with this ability being highest for *Vca*GbpA and lowest for *Lm*LPMO10A.

**Figure 5.**
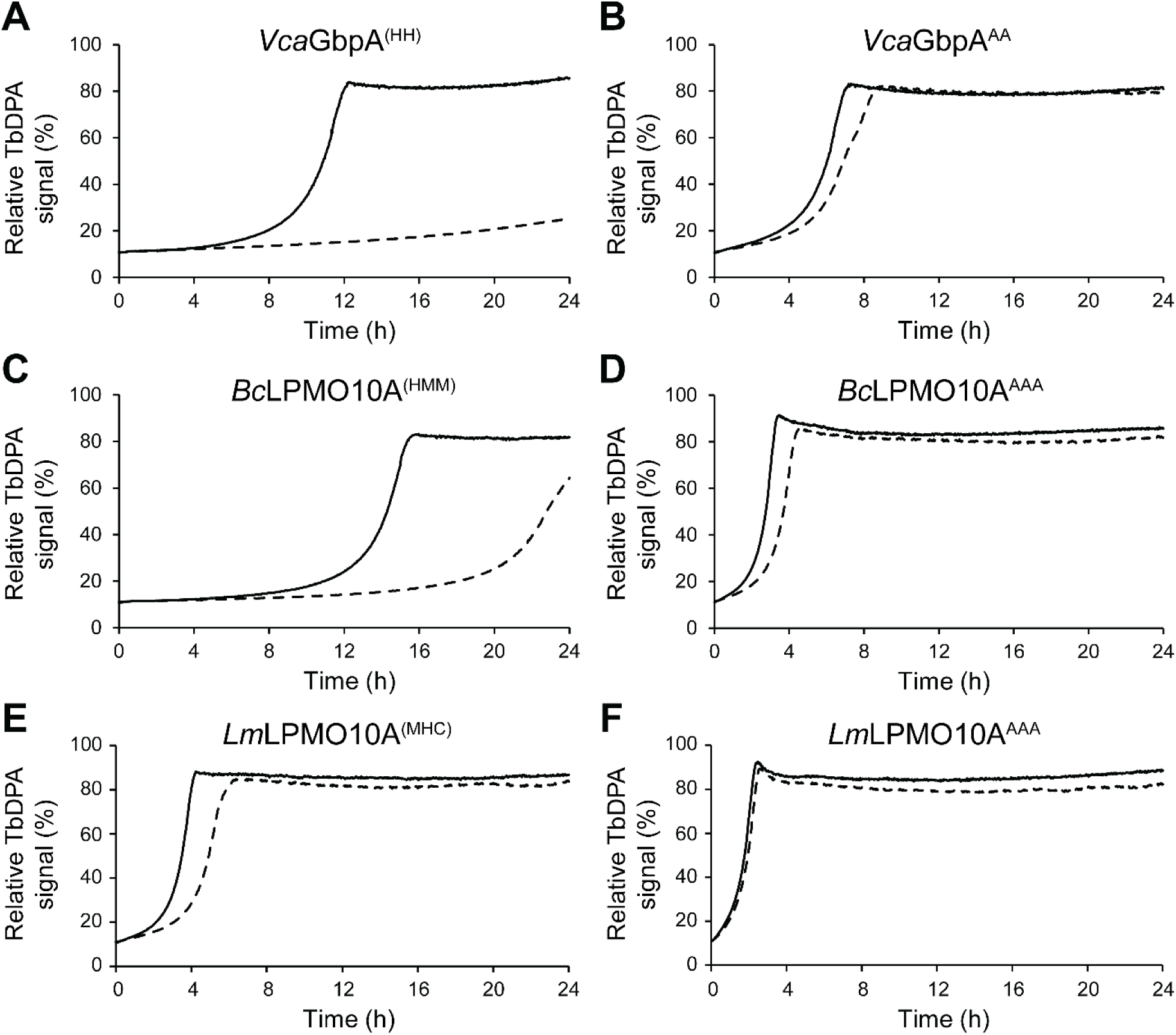
Ascorbic acid depletion in the absence and presence of chitin. The figure shows ascorbic acid oxidation in the presence (dashed line) or absence (solid line) of β-chitin (2 g/L) for wildtype and alanine-substituted variants of *Vca*GbpA (**A&B**), *Bc*LPMO10A (**C&D**), and *Lm*LPMO10 (**E&F**). All reactions contained 1 µM LPMO, 7.5 mM TbCl3, 250 µM DPA, 130 mM sodium acetate buffer at pH 5.6, and were performed in the presence or absence of 1 mM ascorbic acid. The y-axis shows the fluorescence signal from reactions with ascorbic acid relative to the reactions without ascorbic acid. Reactions were carried out in triplicates (n=3), and the mean value is shown for clarity.

Comparison of the wildtype enzymes (Figure 5A,C,E) with variants lacking the copper binding residues in the third domain (Figure 5B,D,F) shows that the mutated variants inactivate more rapidly, both in the absence and presence of substrate. The fact that the stabilizing effect of the substrate is much reduced in the absence of the copper binding site in the third domain underpins the importance of the copper-binding ability of this domain for the stability of the system. Taken together, the results shown in Figure 5 demonstrate that the FnIII domains in all three LPMOs stabilize Cu(I) in the presence of oxygen, and that abolishing this function dramatically affects ascorbic acid oxidation.

The ability to sequester low amounts of reduced copper was assessed by investigating whether the *Bc*FnIII_2 and *Vc*GbpA3 domains could limit oxidation of ascorbic acid in buffer, which is known to be promoted by trace amounts of copper (in the lower nanomolar range, < 50 nM). The results (Figure S11) show that each of these domains effectively reduces the rate of ascorbic acid oxidation, on par with, or even more effectively than EDTA. Thus, these domains are capable of removing trace amounts of copper from the solution, indicative of high affinity for Cu(I).

### Beyond polysaccharide oxidation: The role of multidomain LPMOs in bacterial host interaction and virulence

The multidomain LPMOs studied here share a conserved modular architecture in which a chitin-active N-terminal LPMO domain and one or two C-terminal CBM domains are separated by two lineage-specific domains of previously unknown function. We show that the second internal domain (i.e., the third domain) from Gram-negative bacteria shares the fibronectin III-like fold of Gram-positive bacteria (Figure S5) and that all these domains bind Cu(I) using genus-specific binding motifs (Figure 4). These motifs are only present when the appended catalytic domain is an LPMO domain (Figure S2&3) and lack when the appended catalytic domain is a chitinase (GH18) domain (Figure S4). This reveals that the two central domains of unknown function are not incidental linkers, as has been suggested for the LPMOs of Gram-positive origin [38, 44, 45, 63], but instead constitute dedicated copper-binding modules capable of coordinating Cu(I). While the LPMO domains of these proteins are active on chitin, their conserved multidomain architecture, including a specific Cu(I)-binding module, their expression under host-mimicking conditions, and their demonstrated ability to interact with mucin and host cells [8, 9, 19, 64, 65] point to a role other than chitin degradation. This notion is supported by several studies showing that expression of *Bc*LPMO10A [66, 67], *Ba*LPMO10A (from *B. anthracis*; [68]), *Lm*LPMO10A [45] and *Vc*GbpA [69] is not induced by chitin.

In terms of biological function, *Vc*GbpA has been studied more extensively than the other proteins discussed here. It has been shown that intestinal mucin stimulates production of *Vc*GbpA and, conversely, that *Vc*GbpA stimulates secretion of mucins from intestinal cells, both in a dose-dependent manner [19]. More recent studies show that *Vc*GbpA can trigger a necrotic response in host cells [64] and promote the production of interleukin 8 (IL-8), a pro-inflammatory cytokine, in epithelial cells, thus modulating the host immune response [65]. All in all, it seems clear that *Vc*GbpA has both physical and regulatory interactions with the intestinal epithelium, while there is no obvious role for the polysaccharide-degrading ability of the protein in the gut. In light of the recent findings that the primary intestinal mucin, MUC2, is important for copper uptake and regulation [15, 16] and that copper regulates the host innate immune response against bacterial infections [17], it is conceivable that the ability of the GbpA3 domain to selectively bind Cu(I) has evolved in response to a host environment where copper may be limiting and/or an important regulator of both host and bacterial well-being.

A similar pattern emerges from studies of multidomain LPMOs in the *Bacillus cereus* group. These proteins were first detected in the secretome together with toxins and virulence factors (including mucin-degrading metalloproteases) under conditions unrelated to chitin utilization [66, 68]. Most interestingly, subsequent secretome analyses demonstrated that the multidomain LPMO *Bc*LPMO10A (formerly annotated as chitin-binding protein A, ChbA; gene BC_2798) is strongly enriched under anaerobic conditions, i.e., conditions that mimic the small intestine but are not favorable for LPMO activity, while being markedly repressed in oxic environments. In contrast, the single-domain LPMO from the same organism, *Bc*LPMO10B (gene BC_2827), displayed the opposite trend and was enriched under oxic conditions [67]. More recently, production of *Bc*LPMO10A was shown to be positively regulated by the translocator protein TSPO in *B. cereus*, which is implicated in virulence-related phenotypes [70, 71].

Comparable observations were made in *B. anthracis*. Expression of the homologous four-domain protein *Ba*LPMO10A (GBAA2793) occurred together with major virulence factors under conditions mimicking host infection while being absent during aerobic growth. Likewise, the corresponding single-domain LPMO, *Ba*LPMO10B (GBAA2827), was upregulated under aerobic conditions[68], which seems compatible with a role in polysaccharide degradation. Moreover, transcript levels of *Ba*LPMO10A (GBAA2793) were strongly reduced in an avirulent *B. anthracis* strain (ΔVollum) [72]. Clearly, the available data suggest that multidomain LPMOs in the *Bacillus cereus* group are important for redox adaptation and virulence rather than for degradation of chitin.

In *Listeria monocytogenes*, the multidomain LPMO *Lm*LPMO10A, also known as *Lmo*2467 or chitin-binding protein, was first identified as a virulence factor, as its deletion reduces the bacterial burden in mice [7]. Subsequent sequencing of clinical *Listeria monocytogenes* outbreak isolates in Germany showed that premature stop codons in the LPMO gene are associated with less virulent clinical isolates [73]. *Lm*LPMO10A was shown to oxidatively cleave chitin and act synergistically with chitinases *in vitro*, but its expression is not induced by the presence of chitin, in contrast to the two chitinases from the same strain [45]. Instead, transcription of *lmo2467* is controlled by the redox-responsive regulator SpxA1, which is essential for pathogenesis [74], with deletion of *spxA1* strongly downregulating transcription of *lmo2467* [75]. Interestingly, genes involved in copper handling, including those encoding the regulatory copper(II)-binding protein YcnI [76] and the copper(II) transporter YcnJ [77] show similar regulatory patterns as the LPMO gene [75]. Thus, like *Vc*GbpA and *Bc*LPMO10A, multidomain *Lm*LPMO10A seems to be associated with pathogenicity and processes related to redox conditions and copper availability, rather than chitin utilization.

Here, we show that all these pathogenicity-associated multi-domain LPMOs contain a copper(I)-binding domain next to the LPMO domain, reinforcing an emerging view that copper is a key factor in host–microbe interactions in the intestine. The conserved multidomain arrangement in these proteins appears to provide a structural framework that couples oxidative polysaccharide cleavage by an interfacial enzyme, copper handling, and GlcNAc recognition to host environments with strong redox gradients [78], intricated and physiologically crucial copper management, and host surfaces displaying GlcNAc.

Unlike the multidomain LPMOs described here, all belonging to intestinal pathogens, the lung-associated trimodular LPMO *Pa*CbpD, a well-documented virulence factor [5, 41], lacks a copper-binding FnIII-like domain (Figure 1). Although speculative, this difference could relate to differences between intestinal and lung mucins and in the mucin-dependent regulation of a copper-mediated redox balance [14, 15]. Notably, while both intestinal and airway mucins have been reported to bind Cu(II), Cu(I) binding has only been observed for intestinal mucins and not for airway mucins [15]. This oxidation-state-specific difference supports a model in which auxiliary copper-binding domains have evolved to function in tissues where the presence and bioavailability of Cu(I) are physiologically important.

## Concluding remarks

The combination of literature data, the increasing notion that copper plays a key role in microbial infections in the intestine [80], and the present discoveries strongly suggests that the multidomain virulence-associate LPMOs of intestinal bacteria play a crucial role in managing copper during infection (Figure 6). One key remaining question is the role of the catalytic LPMO domain and its connection with the copper-binding third domain. The copper-binding function of the third domain is only present when such a domain is coupled to an LPMO, so there must be a link. The third domain may interact with the LPMO domain, to protect it from damaging off-pathway reactions, as previously suggested for *Vibrio* GbpA by Zhou et al [21], or perhaps simply to ensure that the LPMO obtains or keeps its copper in an environment where the competition for copper is fierce. The substrate binding studies do support an interaction between the two domains, but, considering that we could not confirm this interaction with SAXS and the varying magnitude of mutational effects on substrate-binding, this interaction may be a side effect rather than the main purpose of the copper affinity of the third domain. In the meantime, the question remains if and why the catalytic activity of the LPMO domain is important and what it acts on.

**Figure 6.**
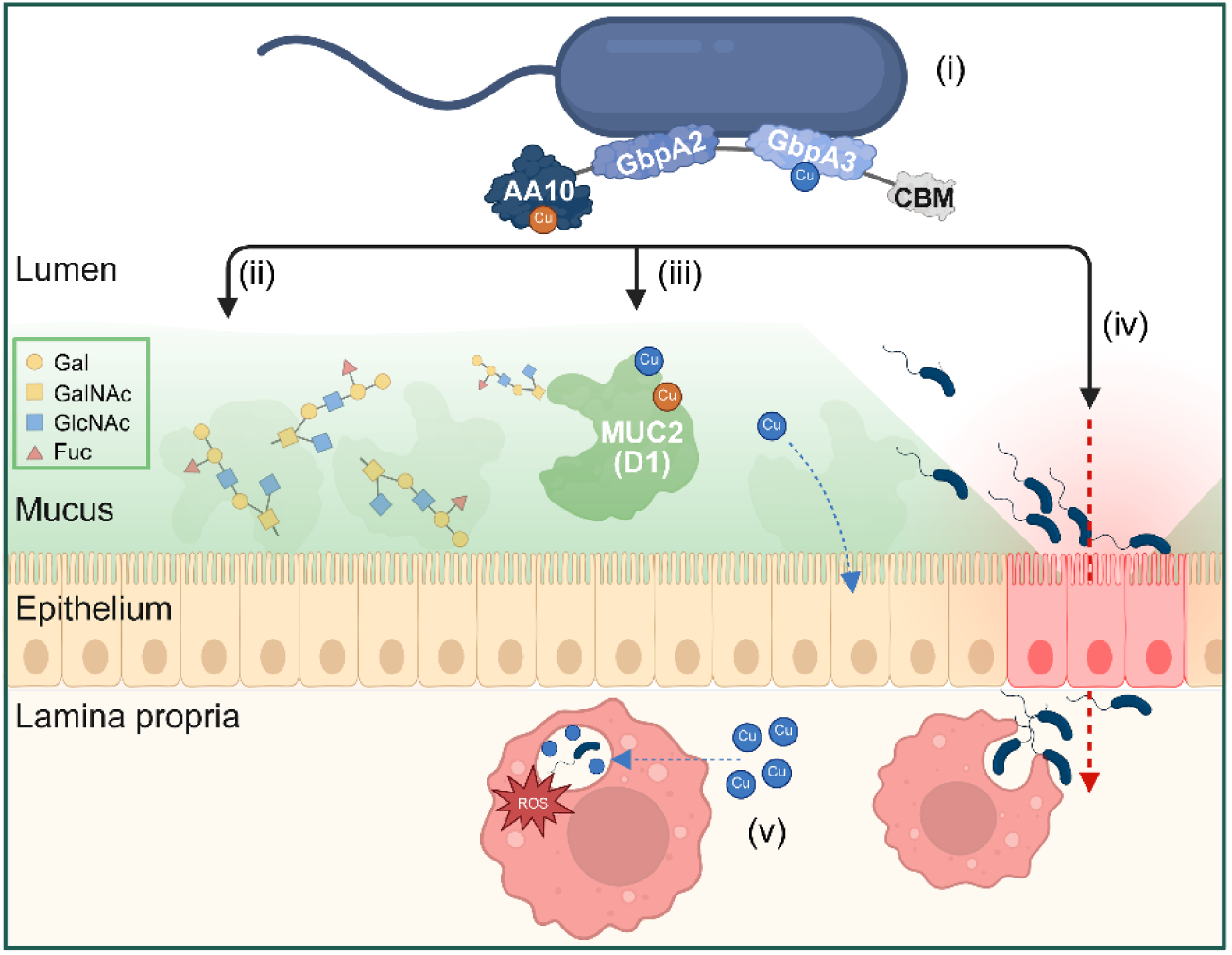
Possible roles of multimodular copper-binding LPMOs during intestinal colonization and infection. This model integrates the present findings with insights from the literature, placing multimodular copper-binding LPMOs in the context of intestinal colonization and infection. In *V. cholerae*, GbpA has been proposed to be displayed on the bacterial surface through interactions mediated by its internal domains (GbpA2–GbpA3), corresponding to the FnIII_1– FnIII_2 domains in *Bc*LPMO10A and *Lm*LPMO10A (i). These proteins may contribute to colonization of the mucus layer through interaction with and, possibly, oxidative cleavage of mucin *O*-glycans (ii). This activity may act in concert with co-regulated virulence factors, such as metalloproteases, capable of degrading mucus components, supporting bacterial colonization and penetration of the mucus layer. Multimodular LPMOs may also compete for copper with host proteins such as MUC2, potentially influencing local copper availability and host–pathogen metal homeostasis (iii). Binding of LPMO domains to epithelial cells may trigger inflammatory responses and promote bacterial adhesion and colonization of the epithelial surface, potentially facilitating bacterial translocation across the epithelium (iv). Following penetration of the epithelium, bacteria will encounter macrophages (v), where they are exposed to antimicrobial conditions including reactive oxygen species (ROS), acidification, and elevated levels of bioavailable copper, all of which challenge bacterial survival [79].

The third domain binds copper with high affinity as demonstrated by its ability to bind trace amounts of copper occurring in common buffers. It is conceivable that these multidomain LPMOs, which have evolved to act in the intestine, in fact are bi-functional proteins in which the primary role of the third domain simply is to sequester copper and thus influence redox and metal homeostasis. Intestinal MUC2 sequesters copper in both redox states [16], and the multidomain LPMOs can do the same (Figure 6). The strong preference of the third domain for Cu(I) fits well with the notion that the intestine is a reducing, anaerobic environment [78]. Since these multidomain intestinal LPMOs are preferentially expressed in the absence of oxygen, [67, 68, 75], i.e., conditions that do not permit LPMO activity, one could envisage that also the role of the LPMO domain primarily, or only, relates to copper homeostasis, at least in infection conditions. These LPMO domains bind both Cu(II) and Cu(I) with high affinity [81, 82].

All in all, it seems clear that a major role of these multidomain intestinal LPMOs is to affect copper levels and copper regulation in the intestine, perhaps competing with MUC2, and affecting one or more steps of the infection process (Figure 6) [15–17, 79, 80, 83]. For the bacterium, copper binding may be beneficial because copper regulates the innate immune response [17]. For invasive pathogens like *Listeria monocytogenes*, copper binding would provide protection against high Cu(I) levels in macrophages. The fact that the modular architecture and copper-binding abilities of these LPMOs are conserved across Gram-positive and Gram-negative intestinal pathogenic bacteria underpins the importance of these intriguing proteins for colonizing the anaerobic, reducing, and mucin-rich intestinal environment in humans and other mammals. The remarkable protective effects obtained after immunizing mice with CbpD from *P. aeruginosa* [41] suggest that the multidomain LPMOs of intestinal pathogens may have potential as vaccine antigens.

## Materials and methods

### Preparation of β-chitin

Commercial β-Chitin (5 mm flakes) was acquired from Glentham Life Science Ltd. (Corsham, UK) and was milled using a PM200 planetary ball mill (Retsch, Haan Germany) with zirconium oxide grinding tools. The milled chitin flakes were sieved and the 200-500 µm fraction was used as model microcrystalline chitin substrate.

### Mutagenesis, cloning, expression, and purification of protein variants

Molecular *in silico* design of genes was performed with SnapGene (Chicago, IL, USA) and protein variants (Table 1) were cloned into the pRSETB vector using the native signal peptide from *Sm*AA10A to ensure correct processing of the LPMO-containing variants and direct the protein to the periplasm. The full-length and truncated wild-type variants were cloned using gene-specific primers and Q5 Hot Start High-Fidelity DNA Polymerase following the manufacturer’s instructions (New England Biolabs, MA, USA). All mutants were subsequently made using the PlatinumTM SuperFi II PCR Master Mix (Thermo Fisher Scientific, MA, USA) following the manufacturer’s instructions for site-directed mutagenesis. PCR reactions contained 0.004 ng template DNA and 0.5 µM of each of the required primers and were performed in 25 µL. 1 µL of the resulting PCR reaction was used to transform *E. coli* One Shot® TOP10 cells (Invitrogen). Single colonies were inoculated and grown overnight in 5 mL BHI (Oxoid) medium supplemented with 100 µg/mL ampicillin (BHI-Amp), at 37°C and 220 rpm. The plasmids were extracted using the NucleoSpin® Plasmid kit (Macherey Nagel, CA, USA) according to the manufacturer’s instructions and the sequence of the mutated gene was verified. Subsequently, plasmids were transformed into *E. coli* One Shot® BL21 Star™ (DE3) cells (New England Biolabs, MA, USA) and single colonies were inoculated and grown overnight in 5 mL BHI-Amp to assess protein expression. Large scale protein expression was performed in 500 mL BHI-Amp in 2 L baffled shake flasks at 37°C for 16-20 h at 225 rpm. Bacterial cells were harvested by centrifugation (6000 rpm) and periplasmic proteins were extracted using an osmotic shock method [84].

Periplasmic extracts were adjusted to 20 mM Tris, pH 8.0, for *Bc*LPMO10A variants including *Bc*FnIII_2, 50 mM Tris (pH 7.5) for *Vca*GbpA variants, and to 50 mM Bis-Tris (pH 6.5) for *Lm*LPMO10A variants prior to loading these solutions onto a 5 mL Q Sepharose fast flow anion-exchange column (Cytiva, MA, USA). Bound proteins were eluted with a 0-500 mM NaCl gradient over 20 column volumes for *Vca*GbpA and *Lm*LPMO10A variants, whilst *Bc*LPMO10A variants were eluted with a 0-200 mM NaCl gradient over 20 column volumes. Fractions containing relevant proteins were identified by SDS-PAGE, pooled and concentrated using Amicon® Ultra centrifugal filters (3 or 10 kDa MWCO depending on the M_w_ of the protein; Millipore, Darmstad, Germany). Final purification was achieved by size-exclusion chromatography using a ProteoSEC Dynamic 16/60 3-70 HR column (Protein Ark, Sheffield, UK) in 50 mM Tris (pH 7.5) and 200 mM NaCl at 1 mL/min. Fractions containing pure protein were pooled and concentrated.

Wildtype and mutated *V. campbellii* GbpA3 (*Vca*GbpA3^(HH)^ and *Vca*GbpA3^AA^) were initially cloned with the *Sm*LPMO10A signal peptide into pRSETB (as for other constructs in this study) and pJB (previously used for inducible periplasmic expression of LPMOs; [85]), but no detectable expression was obtained. As a backup, codon-optimized (for *E. coli*) genes encoding wildtype *Vc*GbpA3 and its H334A/H365A mutant from *V. cholerae* (UniProtKB: Q9KLD5, residues 317–414; 51% sequence identity to *V. campbellii* GbpA3) were purchased as synthesized constructs (cloned into pET22b with a pelB signal peptide GenScript, NJ, USA). A stop codon was introduced upstream of the optional C-terminal His-tag in the pET22b vector to abolish this tag, which could lead to nonspecific copper binding. The pET22b-*Vc*GbpA3 (wildtype and mutant) plasmids were transformed into *E. coli* One Shot® BL21 Star™ (DE3) cells (Invitrogen). Single colonies were inoculated and grown overnight in 5 mL LB medium containing 50 µg/mL ampicillin (LB-Amp) at 37 °C. 2-mL of preculture were inoculated into 400 mL LB-Amp followed by incubation at 30 °C to OD_600_ ≈ 0.6 and, then, induction with IPTG (0.25 mM final concentration). Cultures were then incubated for another ∼16 h at 22 °C and harvested by centrifugation (6,500 rpm). Periplasmic proteins were extracted using the osmotic shock method applied throughout this study.

Periplasmic extracts were adjusted to 30 mM Tris (pH 7.5) and applied to a 5 mL Q Sepharose anion-exchange column (Cytiva, Ma, USA). Bound proteins were eluted with a 0–500 mM NaCl gradient over 60 column volumes. Fractions containing *Vc*GbpA3^(HH)/AA^, identified by SDS–PAGE, were pooled and concentrated using Amicon® Ultra centrifugal filters (3 kDa MWCO; Millipore, Darmstadt, Germany). Final purification was achieved by size-exclusion chromatography on a ProteoSEC Dynamic 16/60 3–70 HR column (Protein Ark, Sheffield, UK) in 50 mM Tris (pH 7.5) and 200 mM NaCl at 1 mL/min. Fractions containing pure protein were pooled and concentrated.

Purified LPMO variants were incubated with a twofold molar excess of CuSO_4_ at room temperature for 30 min to ensure full copper loading. Excess, unbound copper was removed by repeated buffer exchange via dilution and concentration using Amicon Ultra centrifugal filters and 50 mM Tris/HCl buffer (pH 7.5). The cumulative dilution factor exceeded 10^6^, effectively eliminating free copper ions from the samples.

### Chitin-binding assay

The chitin-binding properties of CBM-truncated variants of *Vca*GbpA, *Bc*LPMO10A, and *Lm*LPMO10A were assessed using β-chitin as the substrate. Copper-saturated enzyme variants, including wildtypes and alanine substituted variants, were incubated with 10 g/L (w/v) β-chitin in 50 mM Tris-HCl buffer (pH 7.5). Binding reactions were performed either in the absence or presence of 1 mM ascorbic acid at 22 °C with agitation at 800 rpm for 30 min.

Following incubation, the chitin-bound and unbound enzyme fractions were separated by vacuum filtration using a vacuum manifold (Merck Millipore, Burlington, MA, USA). The filtrate, representing the non-bound enzyme fraction, was collected and the protein content was quantified using the Bradford protein assay (Bio-Rad Laboratories, Hercules, CA, USA). Enzyme binding was determined from the decrease in protein concentration in the supernatant relative to the initial enzyme concentration.

### Small angle X-ray scattering (SAXS)

*Bc*LPMO10A^(HMM)^, *Lm*LPMO10^(MHC)^, *Vca*GbpA^(HH)^ and CBM-truncated *Bc*LPMO10A (i.e., *Bc*LPMO10A-ΔCBM^(HMM^) were expressed and purified as described above. All proteins were copper-saturated as described above. For preparation of the *apo* form, CBM-truncated *Bc*LPMO10A was treated with 10 mM EDTA overnight, followed by buffer exchange using Amicon Ultra centrifugal filters to remove residual chelator and copper ions. Prior to SAXS analysis, all proteins were desalted using PD-10 columns (Cytiva, Marlborough, MA, USA) equilibrated with 50 mM Tris (pH 7.5) supplemented with 250 mM NaCl. SAXS measurements were conducted both in the absence and presence of 1 mM ascorbic acid (AscA). When included, AscA was added immediately prior to sealing the sample and loading it into the SAXS chamber to minimize prolonged exposure. AscA was prepared fresh by dissolving it in the same buffer used for protein desalting.

For all samples, scattering intensities, *I(q)*, were recorded as a function of the scattering vector *q*, defined as:

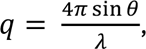

where *λ* is the X-ray wavelength and *θ* is half the scattering angle.

Measurements of *Bc*LPMO10A^(HMM)^, *Lm*LPMO10A^(MHC)^, and *Vca*GbpA^(HH)^ were performed in 1.5 mm thick glass capillaries (Hilgenberg, Malsfeld, Germany, Catalogue no. 4007,615) with a Xenocs Xeuss 3.0 SAXS instrument (Xenocs, Grenoble, France) equipped with a GeniX3D Cu Kα X-ray source (wavelength = 0.154 nm). Scattering intensities were detected using a Pilatus 300K pixel detector (Dectris, Switzerland) at a sample-to-detector distance of 1200 mm, yielding a usable q-range of 0.006 – 0.4 Å^−1^.

Buffer scattering profiles were recorded under identical conditions and subtracted from the corresponding protein measurements. Protein concentrations were 8 mg/mL for *Lm*LPMO10^(MHC)^, and *Vca*GbpA^(HH)^, and 2 mg/mL for *Bc*LPMO10A^(HMM)^. Monodispersity of the samples was confirmed by Guinier analysis. Data processing was done using XSACT 2.4 software (Xenocs, Grenoble, France) in combination with in-house scripts in MATLAB (The MathWorks, Inc., Natick, MA, USA).

For CBM-truncated *Bc*LPMO10A variant, SAXS data were collected at the BM29 beamline at the European Synchrotron Radiation Facility (ESRF, Grenoble, France) using a flow-through capillary cell and an X-ray wavelength of 0.1 nm. Measurements were conducted at 20 °C, with data recorded in ten successive 1 s frames, yielding a usable q-range of 0.0045–0.52 Å^−1^. Frames exhibiting radiation damage were excluded, and the remaining frames were averaged for further analysis. Buffer scattering data were acquired using the same experimental setup and subtracted from the corresponding sample data. All protein samples, in both the absence and presence of reductant, were measured at concentrations of 0.5, 1, 2, 4, and 8 mg/mL. The scattering data were extrapolated to zero protein concentration using *PRIMUS* from the *ATSAS* software package. Pair-distance distribution functions, *P(r)* were calculated using GNOM. Conformation models were obtained by fitting the data with MultiFoXS, using an AlphaFold [86] prediction model as the starting model. Linker regions were defined as flexible. MultiFoXS generated 10,000 different conformations. The final model corresponds to the best-scoring two-state ensemble.

### Copper binding using BCS

To assess copper binding by *Vc*GbpA and *Bc*FnIII_2 (wildtype and alanine-substituted variant), 4 µM protein was incubated with 8 µM Cu(II)SO_4_ in 50 mM sodium phosphate buffer (pH 6.0) for 10 min at room temperature, with or without 20 µM ascorbic acid. Proteins were removed using 3 kDa MWCO microcentrifugal filters (VWR, catalog 82031-346) for 5 min at 10,000 × g, and 100 µL of the filtrate was transferred to black 96-well plates (Thermo Fisher). Residual (unbound) copper was quantified using bathocuproine disulfonate (BCS), a Cu(I)-specific probe whose fluorescence decreases upon copper binding [87]. Each filtrate was mixed 1:1 with 40 µM BCS in 50 mM sodium phosphate buffer (pH 6.0) containing 40 µM ascorbic acid and incubated 10 min at room temperature in a Varioskan LUX plate reader. Fluorescence (λ_ex_/λ_em_ = 290/325 nm) was monitored every minute for 10 min to confirm signal stability. Free copper concentrations were calculated from the average fluorescence intensity over the 10 min measurement period using a standard curve prepared with CuSO_4_ (0-8 µM) treated identically to the samples.

### Density functional theory (DFT) calculations

The initial structural models of the FnIII-2 domains of *Lm*LPMO10A and *Bc*LPMO10A and of GbpA3 from *Vc*GbpA were obtained using AlphaFold3 [46] and contained 88, 95, and 98, residues respectively, one Cu(II) ion per protein, and one Ca(II) ion for the two FnIII_2 domains. Hydrogen atoms were added using pdb4amber [88]. Protonation states were assigned at pH 4 based on p*K*_a_ values predicted by PROPKA 3.5 [89]. Geometry optimization was performed using the GFN2-xTB semiempirical tight-binding method [50] as implemented in ORCA 6.1 [90], with the analytical linearized Poisson–Boltzmann (ALPB) implicit solvation model for water [91].

### Electron paramagnetic resonance spectroscopy (EPR)

Continuous-wave X-band (∼9.47 GHz) EPR spectra were acquired using a Bruker Magnettech ESR5000 (Bruker, Billerica, MA, USA). Measurements were performed in custom-made quartz EPR tubes (4 mm outer diameter). Samples were frozen in liquid nitrogen (77 K), and spectra were recorded at 80 K with a 60 s sweep time, 100 kHz modulation frequency, 1 mT modulation amplitude, and 10 mW microwave power. Spectral analysis was carried out in MATLAB using the EasySpin package (version 6.0.9) [92]. Prior to setting up the experiments, the proteins were copper saturated as described above; the final protein concentration used in the EPR measurements was 300 µM. The cumulative dilution factor achieved during this process exceeded 10^6^, effectively eliminating free copper ions from the samples.

### Ascorbic acid oxidation – fluorescence

Ascorbic acid oxidation in the absence of chitin particles can be measured directly by recording A_255_, but the presence of chitin particles interferes with this absorption signal. To assess oxidation of ascorbic acid in presence of chitin particles, we developed a method based on the known ability of ascorbic acid to quench the luminescence signal of a terbium-2,6-dipicolinic acid (TbDPA) complex [58]. Luminescent lanthanide complexes such as TbDPA exhibit long excited-state lifetimes and large Stokes shifts, leading to a substantial separation between excitation and emission wavelengths. For TbDPA, excitation occurs at 270 nm and emission at 545 nm. Upon excitation, DPA absorbs a photon and the corresponding energy absorbed is transferred from DPA to the Tb^3+^ ion resulting in luminescence. In presence of ascorbic acid, the luminescence signal of TbDPA is quenched in proportion to the amount of ascorbic acid present, an effect that likely is caused by binding of ascorbic acid to the TbDPA complex and interference with the energy transfer from DPA to Tb^3+^ [58].

The fluorescence signal of the TbDPA complex is stable between pH 5 and 7 and shows a maximum intensity at pH 6 [93]. We found that 130 mM sodium acetate pH 5.6 resulted in the strongest fluorescence signal in our reaction setup. The relatively high acetate concentration likely reduces water accessibility to the coordination shell of the TbDPA complex resulting in a stronger fluorescence signal [93].

Ascorbic acid oxidation assays were performed in black 96-well fluorescence microtiter plates using a Varioskan Lux plate reader (Thermo Fisher Scientific). 1 µM LPMO was pre-incubated with 2 g/L (w/v) β-chitin (Glentham Scientific, 200-500 µm fraction), 7.5 mM TbCl_3_, 250 µM DPA in 130 mM sodium acetate buffer (pH 5.6), at 30°C for 5 minutes. The reactions were initiated by addition of ascorbic acid to a final concentration of 1 mM and the fluorescence signal was measured (λ_ex_ = 270 nm, λ_em_ = 545 nm) every 2. minute over 24 hours at 30 °C without sealing the plate. Each reaction was performed with and without substrate, as well as with and without ascorbic acid, with three independent replicates for each reaction. Ascorbic acid was prepared in TraceSelect water (Honeywell, Seelze, Germany) and its oxidation is reported as the relative TbDPA signal (%), which represents the fluorescence signal from the reaction with ascorbic acid divided by the signal from the reaction without ascorbic acid, multiplied by 100.

### Ascorbic acid oxidation – absorbance

Ascorbic acid oxidation in absence of substrate was measured directly by monitoring the decrease in A_255_ using UV-transparent 96-well microtiter plates and a Varioskan Lux plate reader (Thermo Fisher Scientific). 50 µM stock solutions of wildtype and alanine-substituted*Bc*FnIII_2 and *Vc*GbpA3 were prepared in 50 mM sodium phosphate (pH 7.5). Control reactions were prepared similarly and included 50 mM sodium phosphate (pH 7.5) with or without EDTA or CuSO_4_. The reaction mixtures contained 1.33 µM LPMO, 1.33 µM EDTA or 1.33 µM Cu SO_4_. After 5 min preincubation at 37 °C, ascorbic acid was added to a final concentration of 1 mM, followed by 10 s of shaking prior to measuring A_255_ every 30 s for 4 h. The stock solution of ascorbic acid was prepared freshly in TraceSelect water. Reactions were performed as three independent replicates.

## Acknowledgements

The work was supported by funding from the European Research Council (ERC) through a Synergy Grant (Grant No. 856446). The computations were performed on resources provided by Sigma2 - the National Infrastructure for High-Performance Computing and Data Storage in Norway, project numbers NN1003K and NS1003K. We acknowledge the European Synchrotron Radiation Facility (ESRF) for provision of synchrotron radiation facilities under proposal ID MX-2788 and on beamline BM29. We thank Hayden Fisher for assistance and support during the beamtime. We also thank Ingeborg Langeby Korsane for her contribution to developing the TbDPA fluorescence-based method for ascorbic acid monitoring and Geir Mathiesen for helpful discussions.

## Author contributions

**EGK, HB, VGHE,** and **ZF** designed the research. **EGK, HB, SER, HVS, AR, YZ, ÅKR,** and **ZF** performed the research. **EGK, HB, SER, HVS, AR, YZ, UK, ÅKR, VGHE**, and **ZF** analyzed the data. **EGK, VGHE** and **ZF** wrote the first draft of the manuscript and were involved in finalizing the paper for submission.

