## Supplementary information for "Conserved copper(I)-binding auxiliary domains link virulence-associated LPMOs to copper homeostasis in the intestinal environment"

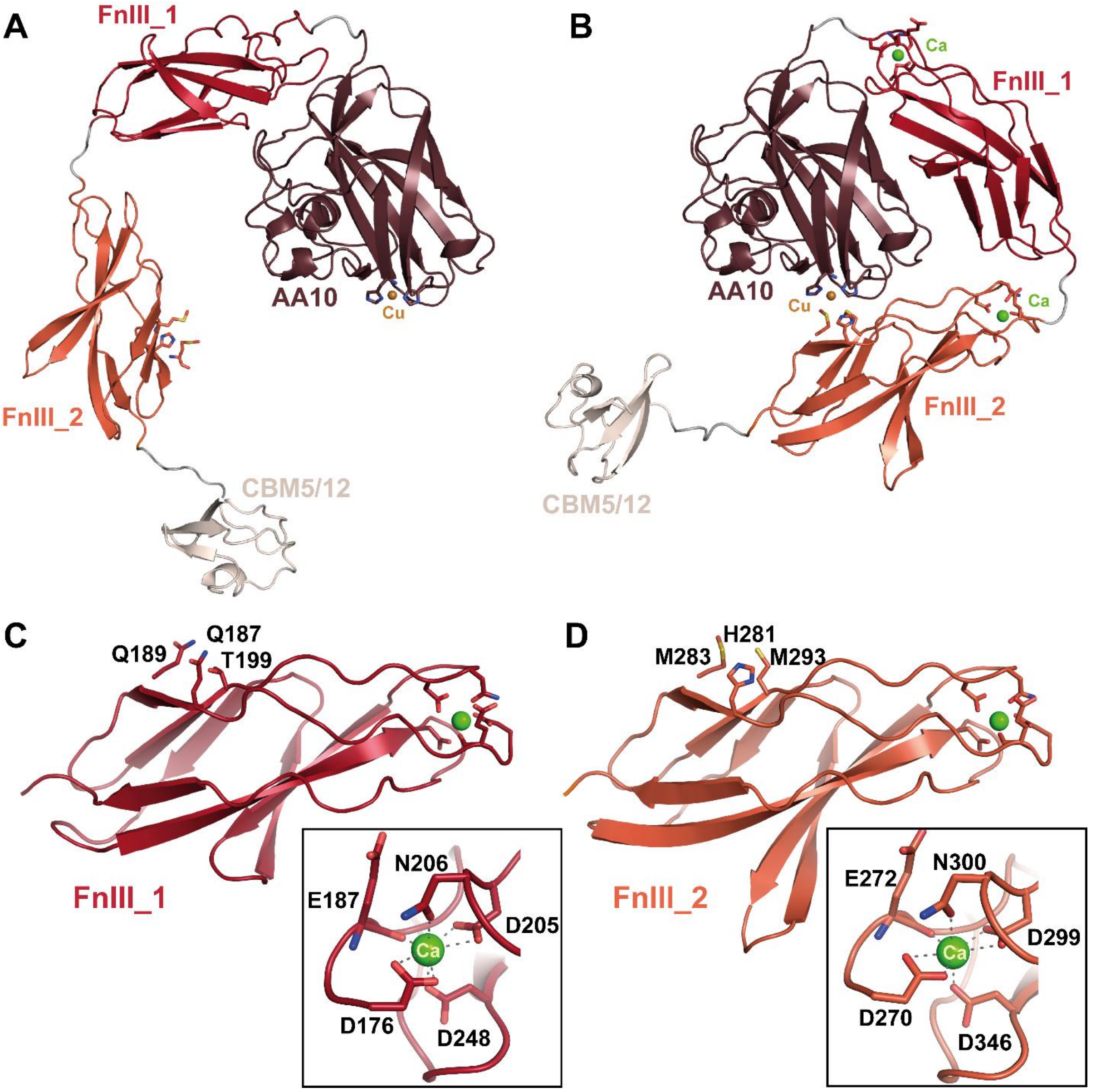


**Figure S1. Predicted calcium-associated conformational changes in *Bc*LPMO10A.** The AlphaFold3-predicted structure of copper-bound *Bc*LPMO10A (A) in the absence of calcium shows an extended conformation comprising four domains: AA10 (catalytic domain), FnIII_1 and FnIII_2 (fibronectin type III–like domains), and CBM5/12 (C-terminal chitin-binding domain). In this model, no contact is apparent between the AA10 and the FnIII_2 domains, in contrast to the (copper-dependent) domain proximity observed in *Vca*GbpA and *Lm*LPMO10A (Figure 2). On the other hand, when including two calcium ions in the prediction, that bind to the FnIII-like domains, the predicted structure adopts a conformation that brings the AA10 and FnIII_2 domain in close proximity. In this predicted arrangement, the catalytic copper coordinated by the AA10 histidine brace is positioned near the conserved HMM motif in the FnIII_2 domain. Structural comparison shows that the two FnIII-like domains of *Bc*LPMO10A are overall similar (C–D), with the HMM motif only present in FnIII_2. In both FnIII domains, $Ca^{2+}$ is predicted to adopt a seven-coordinate, pentagonal bipyramidal geometry, consistent with common calcium-binding motifs such as EF-hands[1].

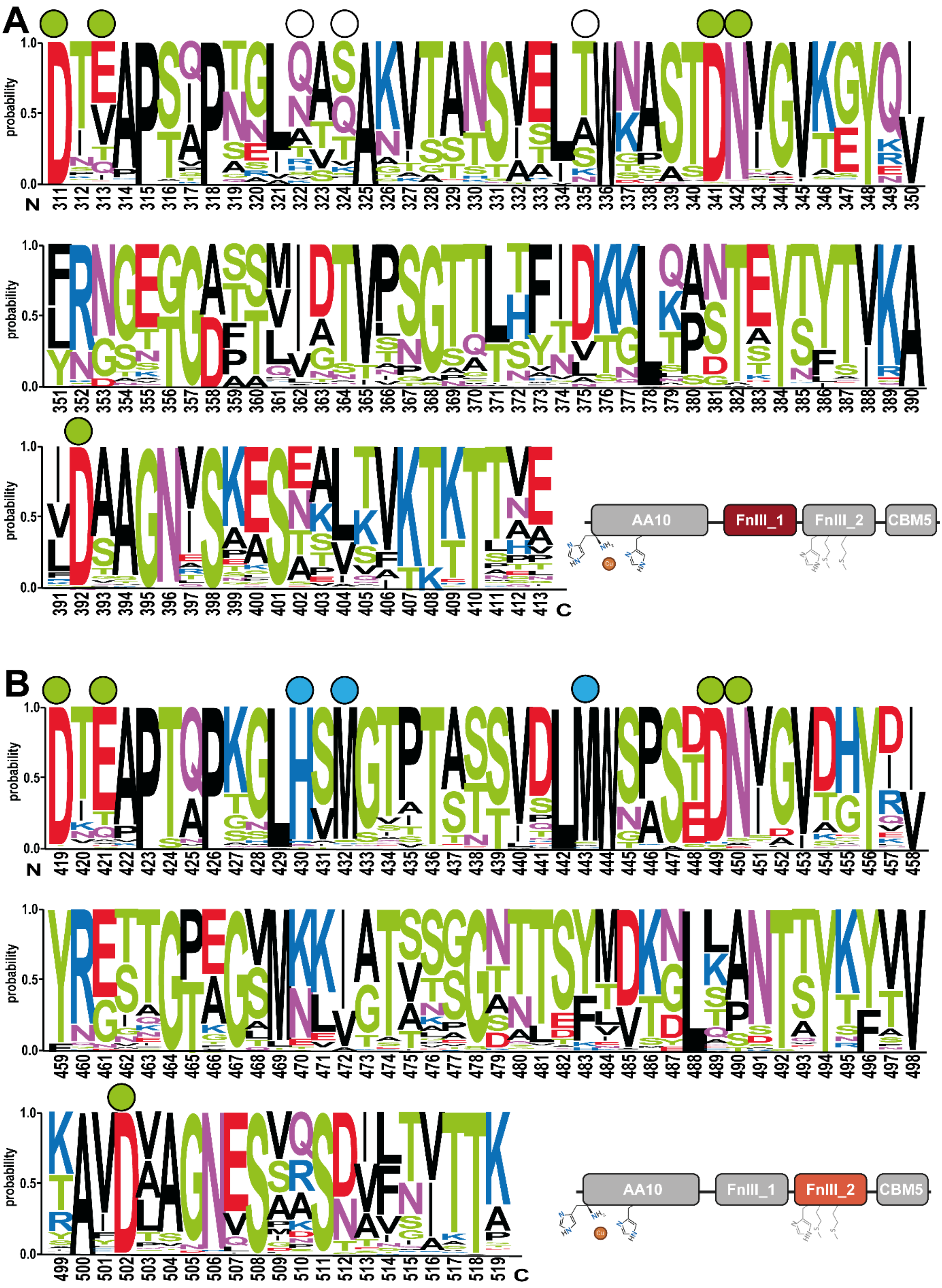


**Figure S2. Sequence conservation in the two internal FnIII-like domains of *Bc*LPMO10A-like proteins.** The figure shows a WebLogo analysis of 590 *Bc*LPMO10A-like sequences. Panels A and B show conservation patterns in the FnIII_1 and FnIII_2 domain, respectively. Green circles indicate residues comprising the putative $Ca^{2+}$-binding site, which is highly conserved in both FnIII domains across *Bc*LPMO10A-like LPMOs sharing this modular architecture (AA10-FnIII_1-FnIII_2-CBM5/12). Blue circles mark residues involved in copper binding in the FnIII_2 domain, whereas open circles denote the corresponding positions in the FnIII_1 domain. The WebLogos show that the putatively copper-binding residues are conserved in the FnIII_2 domain (panel B; HMM motif; His281, Met283 and Met293 in *Bc*LPMO10A) but are absent in the first FnIII domain (panel A).

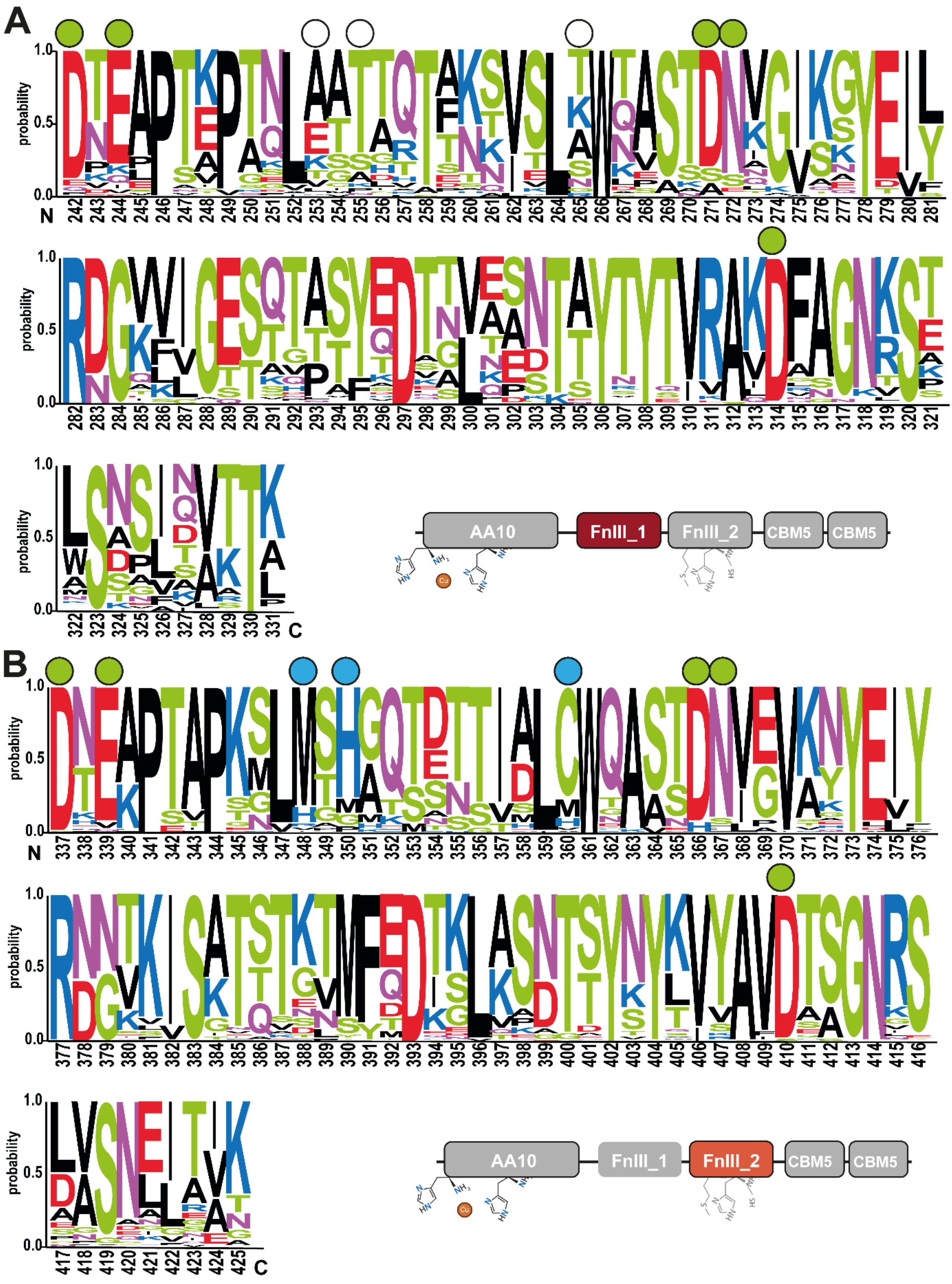


**Figure S3. WebLogo analysis of FnIII domains in 93 *Lm*LPMO10A-like sequences.** Panels A and B show FnIII_1 and FnIII_2, respectively. Green circles mark the conserved putative $Ca^{2+}$-binding site in both domains. Blue circles indicate residues involved in copper binding in the FnIII_2 domain (MHC motif; Met279, His281, Cys291 in *Lm*LPMO10A). The corresponding positions in the FnIII_1 domain are labeled with open circles and lack conserved copper-binding residues.

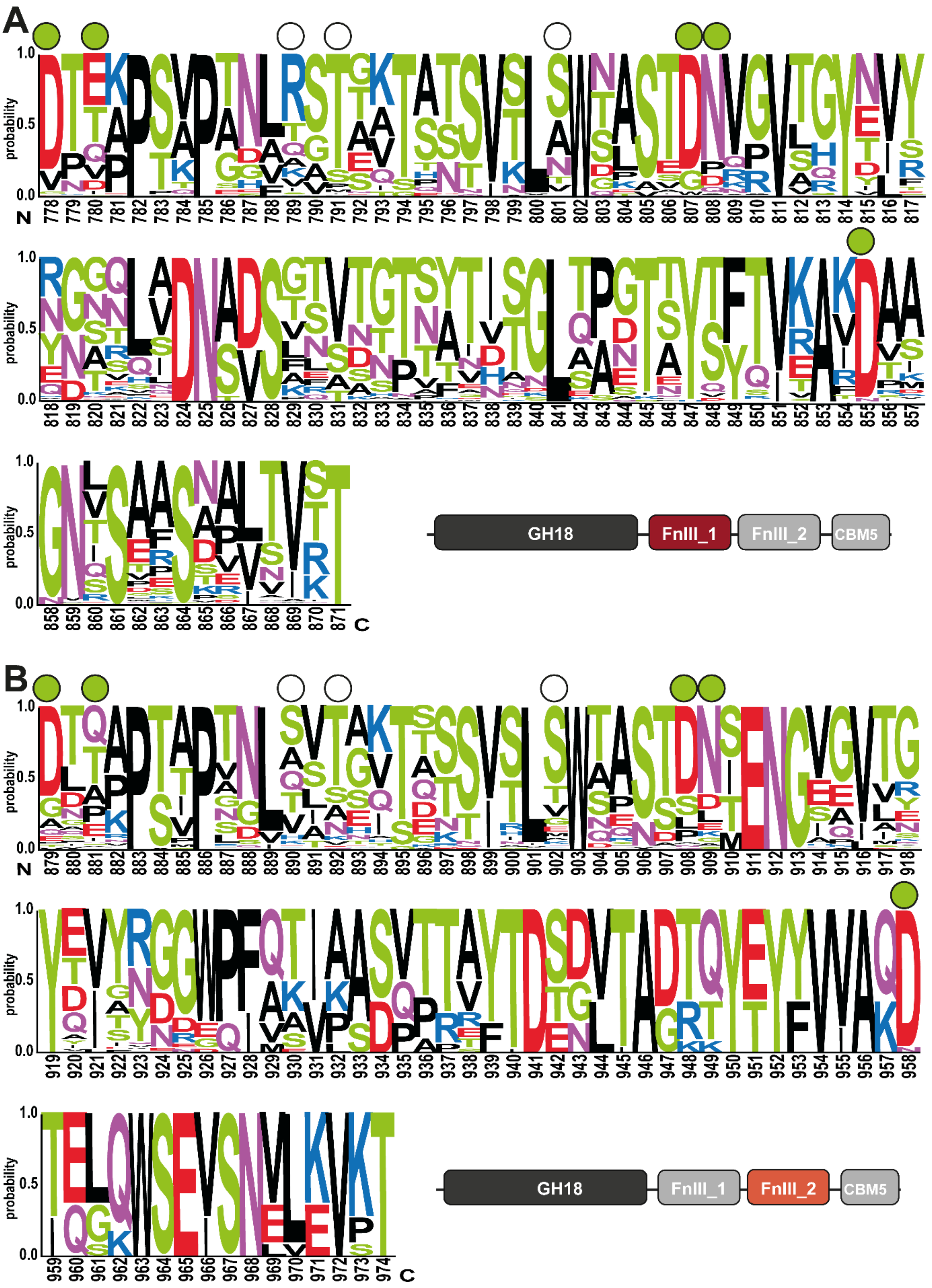


**Figure S4. WebLogo analysis of FnIII domains in 155 GH18 chitinases with a GH18–FnIII_1–FnIII_2–CBM5/12 architecture.** Panels A and B show FnIII_1 and FnIII_2, respectively. Green circles indicate residues at positions analogous to putative $Ca^{2+}$-binding sites, while open circles mark positions corresponding to putative copper-binding sites in LPMO-associated FnIII domains (Figures S2 & S3). Clearly, these chitinase-associated FnIII domains do not show conserved residues putatively involved in copper-binding, whereas residues putatively involved in calcium binding are less well conserved compared to LPMO-associated FnIII domains.

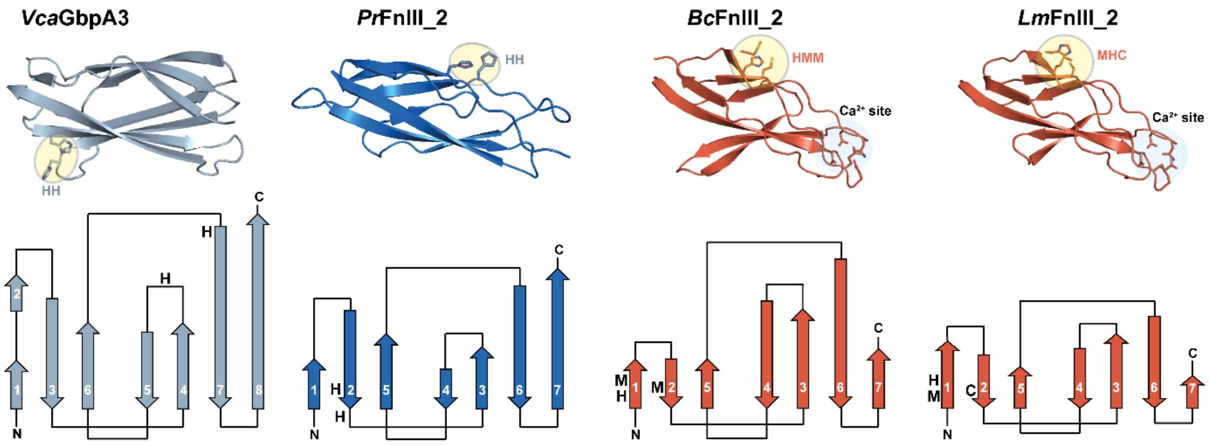


**Figure S5. Structural comparison of third domains in multidomain LPMOs.** The upper row shows the structurally aligned third domains in *Vibrio campbellii* (GbpA3) and the *Bacillus cereus* and *Listeria monocytogenes* LPMOs (FnIII_2), highlighting the relative positions of the putative copper-binding residues. The analysis includes *Pr*FnIII_2 from *Pseudoalteromonas rubra*, a Gram-negative species with an LPMO exhibiting a mixed domain architecture that combines a GbpA2 domain with a FnIII_2 domain (Figure 1). Although the putative copper-binding residues are conserved between *Vca*GbpA3 and *Pr*FnIII_2, the relative position of the copper site within the domain differs, and *Pr*FnIII_2 possesses a predicted $Ca^{2+}$-binding site that is absent in *Vca*GbpA3. The topology diagrams in the lower row show that all third domains share the same overall β-sheet topology, with an identical arrangement of strands, despite variations in strand and loop length.

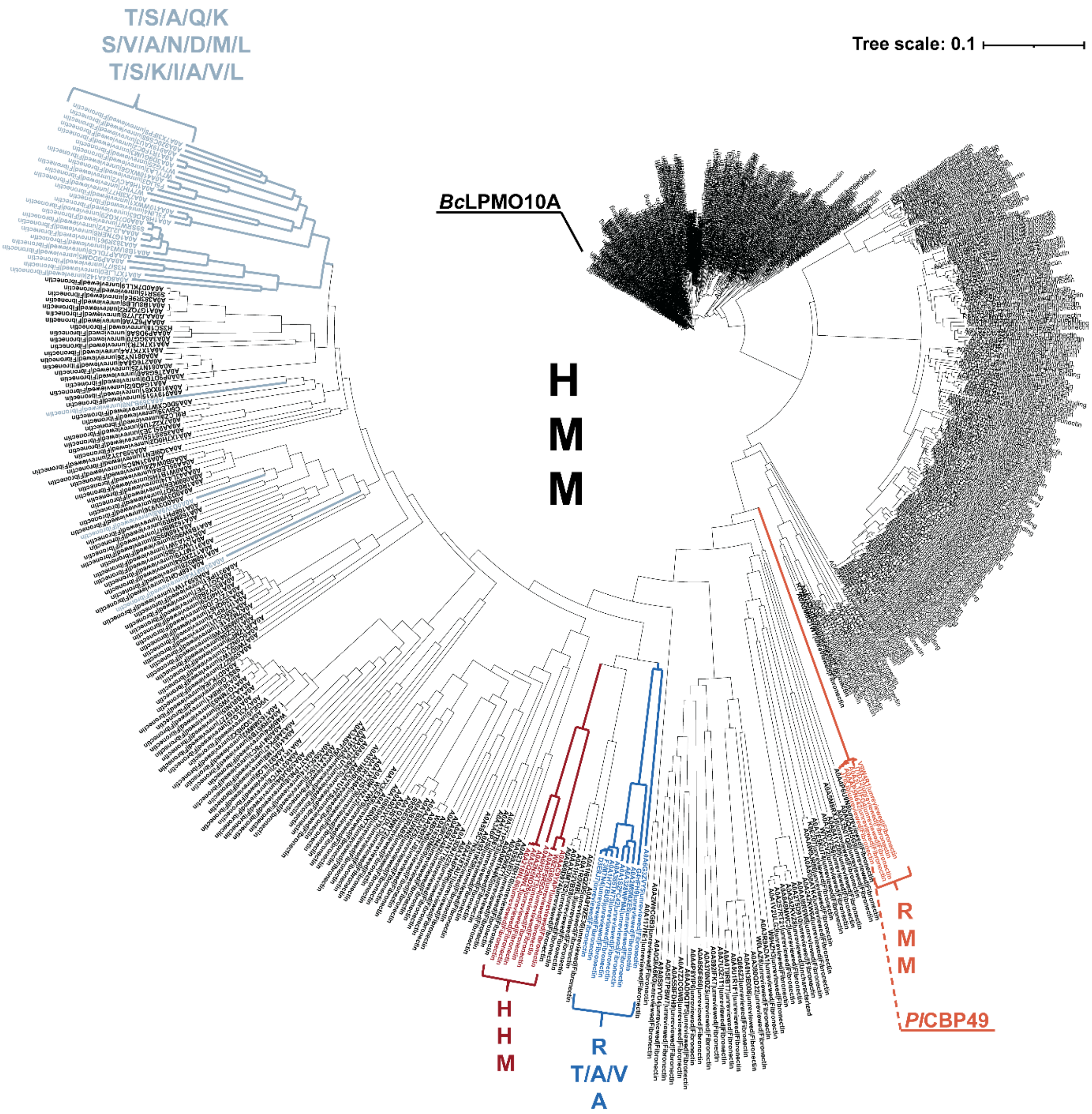


**Figure S6. Phylogenetic tree of 590 sequences sharing the same modular architecture as *Bc*LPMO10A (AA10–FnIII_1–FnIII_2–CBM5/12).** Sequences labeled in black contain the HMM motif in the FnIII_2 domain. The dominant genera represented in the tree are *Bacillus* and *Paenibacillus*, with additional sequences from *Enterococcus*, *Lysinibacillus*, *Pseudomonas*, and *Yersinia*. A subset of 52 sequences, predominantly (>95%) from the genus *Paenibacillus*, display variation in this region, most commonly a loss of the motif. These account for ~30% of the *Paenibacillus* sequences in the tree and appear in various colors (other than black). The phylogenetic positions of *Bc*LPMO10A and *Pl*CBP49 discussed in this study (with *Bc*LPMO10A experimentally characterized) are highlighted in the tree. Figure S2 shows WebLogos representing the FnIII domains of these 590 proteins.

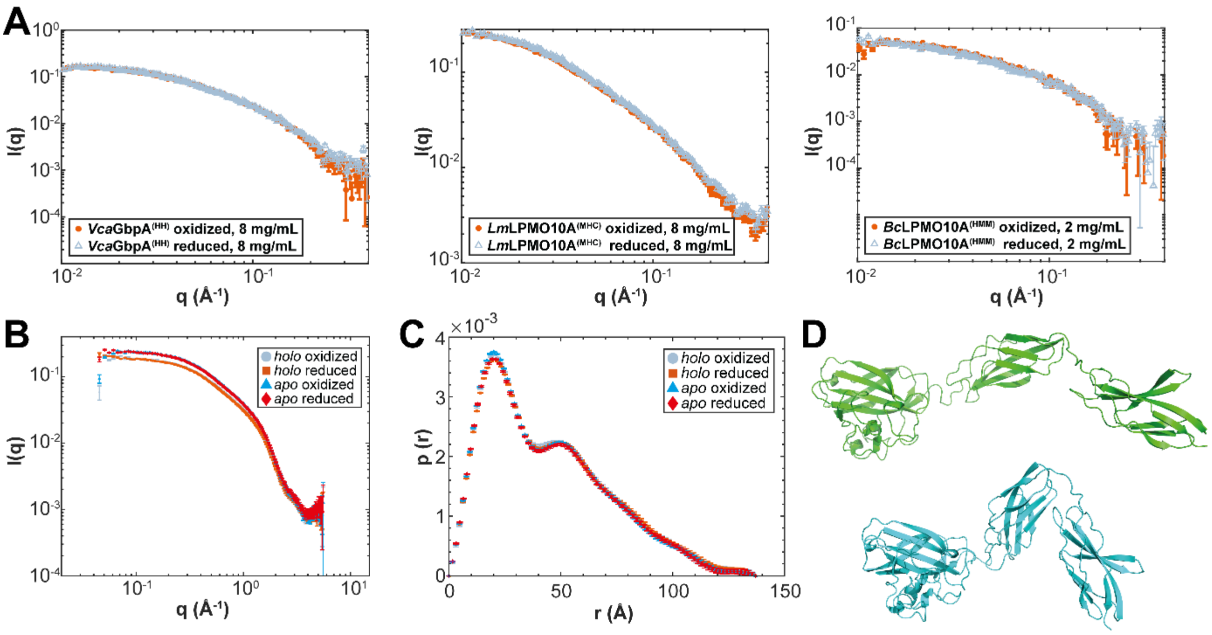


**Figure S7. SAXS analysis.** Panel A shows SAXS scattering profiles for full-length *holo Vca*GbpA(HH), *Lm*LPMO10A(MHC), and *Bc*LPMO10A(HMM) in their oxidized and reduced states. Radius of gyration (*R*g) values derived from the SAXS data were 36.0 ± 0.7 Å and 36.6 ± 0.8 Å for oxidized and reduced *Vca*GbpA(HH), respectively, in agreement with previously reported values for the extended GbpA conformation [2]. Similarly, the *R*g values of *Lm*LPMO10A(MHC) (47.9 ± 0.7 Å oxidized; 47.7 ± 0.8 Å reduced) and *Bc*LPMO10A(HMM) (38.0 ± 1.5 Å oxidized; 41.1 ± 2.0 Å reduced) show only minor redox-dependent differences and are consistent with predominantly extended conformations. Panel B shows SAXS data for *apo*- and *holo*-*Bc*LPMO10A-ΔCBM(HMM) collected under oxidizing and reducing conditions. The scattering profiles are highly similar under all conditions, with only a small offset in absolute intensity between the *apo* and *holo* proteins, consistent with a minor difference in protein concentration. Panel C displays pair-distance distribution functions, P(r), derived from the SAXS data in panel B. The distributions are nearly identical for all samples and show a maximum at low r values (~20 Å) together with a pronounced tail extending to larger distances, consistent with an elongated multidomain architecture. The calculated radius of gyration (*R*g) is 34.5–35 Å under all conditions. SAXS data were fitted using MultiFoXS [3] with models consisting of either a single conformation or ensembles containing up to five conformations. Increasing the ensemble size beyond two conformations resulted in only negligible improvements in fit quality. Panel D shows a two-conformation ensemble model of reduced *holo*-*Bc*LPMO10A-ΔCBM(HMM) generated with MultiFoXS. The upper and lower conformations contribute weights of 0.65 and 0.35, respectively, yielding a good fit to the experimental data ($\chi^2$ = 2.3). Both conformations adopt extended arrangements, although the second is slightly more compact. Neither model indicates interactions between domains 1 and 3.

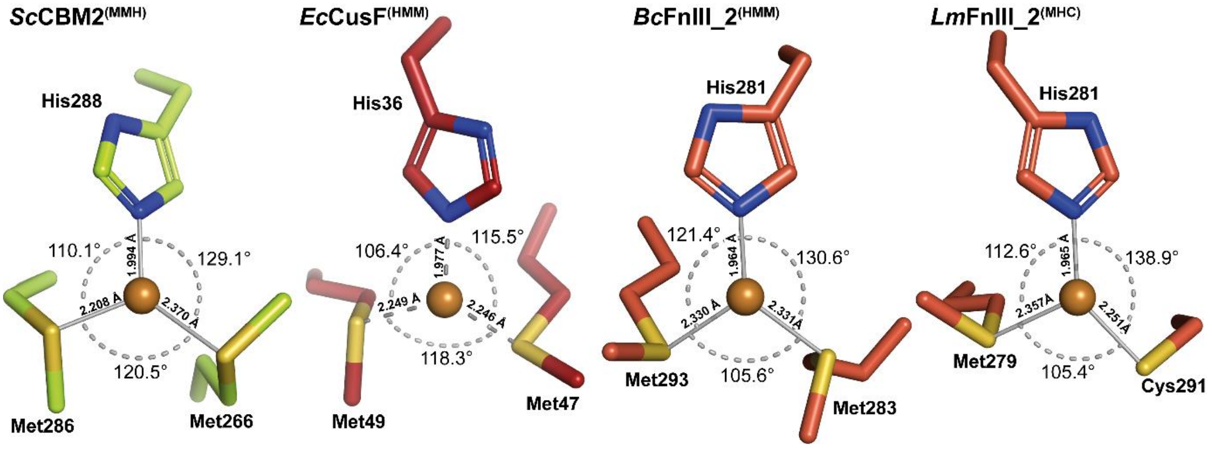


**Figure S8. Geometry-optimized Cu(I) coordination in FnIII_2 domains.** The Cu(I)-binding sites in *Bc*FnIII_2(HMM) and *Lm*FnIII_2(MHC) were geometry optimized to characterize their coordination environments. In both models, Cu(I) adopts a distorted trigonal planar geometry. In *Lm*FnIII_2(MHC), the cysteine residue was modeled in its deprotonated thiolate form, consistent with sulfur coordination to Cu(I). For comparison, the previously published optimized Cu(I) binding sites in *Sc*CBM2(MMH) [4] (see Figure 4 for binding data) and the copper-trafficking protein CusF from *Escherichia coli* (PDB: 2VB2)[5] are also shown. Although sidechain conformations differ, the overall coordination geometries are highly conserved. Pairwise fitting of the coordinating atoms to the CusF crystal structure yielded RMSD values of 0.214 Å for *Lm*FnIII_2(MHC), and 0.219 Å for *Bc*FnIII_2(HMM), indicating close geometric agreement with the experimentally determined CusF Cu(I) site.

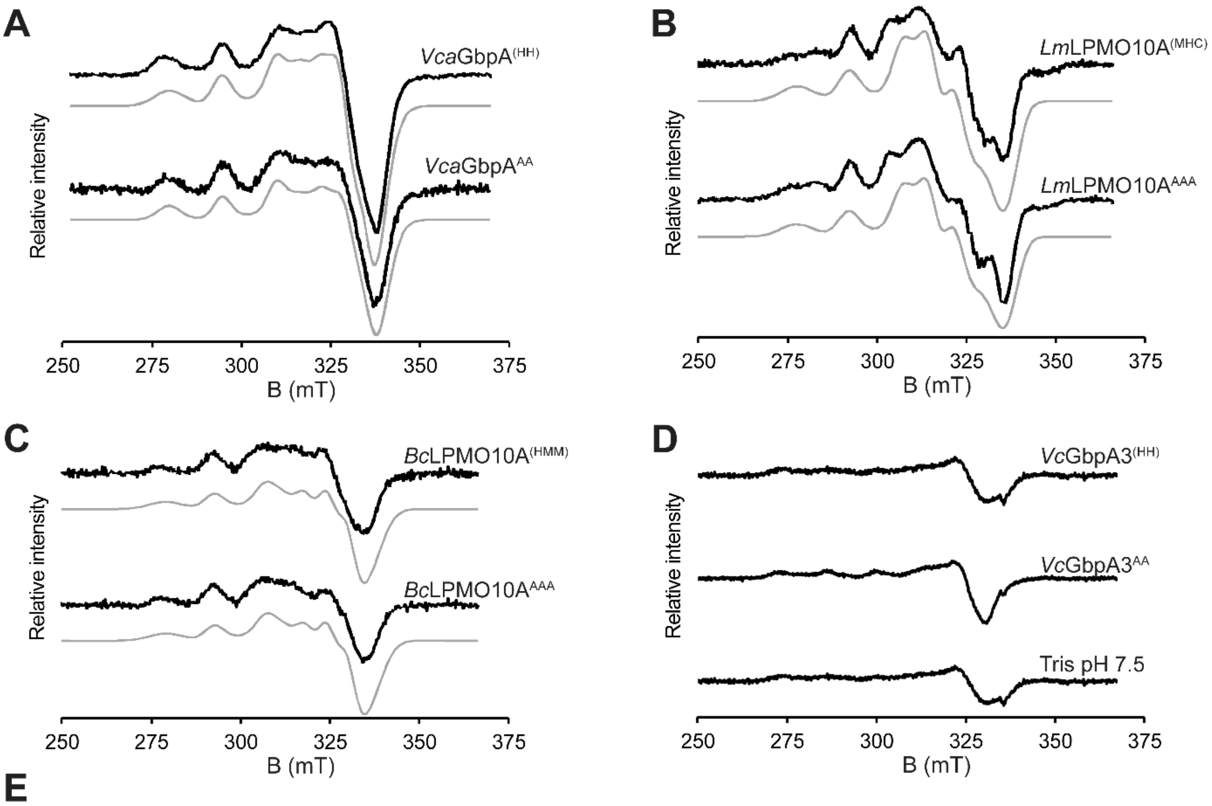


**E**

| Spin Hamiltonians | *Vca*GbpA(HH) | *Vca*GbpA$^{AA}$ | *Bc*LPMO10A(HMM) | *Bc*LPMO10A$^{AAA}$ | *Lm*LPMO10(MHC) | *Lm*LPMO10$^{AAA}$ |
|---|---|---|---|---|---|---|
| $g_z$ | 2.244 | 2.246 | 2.244 | 2.244 | 2.245 | 2.242 |
| $g_y$ | 2.034 | 2.069 | 2.077 | 2.077 | 2.010 | 2.027 |
| $g_x$ | 2.069 | 2.044 | 2.034 | 2.034 | 2.129 | 2.126 |
| $A_z$ | 475 | 470 | 440 | 440 | 470 | 490 |
| $A_y$ | 90 | 140 | 170 | 170 | 30 | 80 |
| $A_x$ | 150 | 130 | 125 | 125 | 20 | 90 |

**Figure S9. Continuous-wave X-band EPR spectroscopy.** Spectra were recorded using 300 μM protein solutions in 50 mM Tris-HCl (pH 7.5). Samples were flash-frozen in liquid nitrogen and analyzed at 80 K. Data were collected using a sweep time of 60 s, a modulation frequency of 100 kHz, a modulation amplitude of 1 mT, and a microwave power of 10 mW. Spectral simulations were performed using the EasySpin 6.0.9 package from MATLAB. The panels A-C show spectra for *Vca*GbpA(HH)/AA, *Lm*LPMO10A(MHC)/AAA, and *Bc*LPMO10A(HMM)/AAA, respectively. The experimental spectra are shown in black, with the simulated spectra shown as grey lines below. Panel D shows the experimental EPR spectra of *Vc*GbpA3(HH)/AA, showing no evidence of Cu(II) binding. The spin Hamiltonian parameters derived from spectral simulations in A-C are listed in panel E.

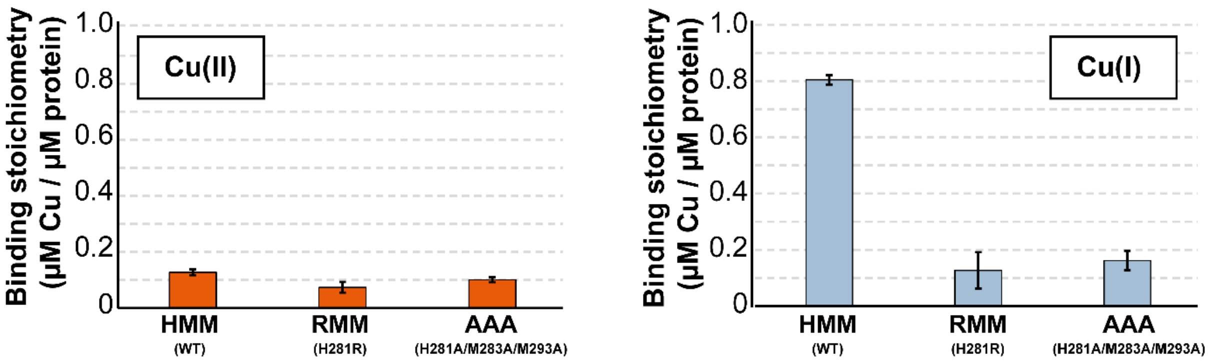


**Figure S10. Copper-binding by *Bc*FnIII_2 variants assessed using the BCS assay.** Copper binding by wild-type *Bc*FnIII_2$^{(HMM)}$ and mutants in which the copper-binding motif had been mutated to RMM or AAA was assessed under oxidizing and reducing conditions using ascorbic acid for reduction [generation of Cu(I)] and the bathocuproine disulfonate (BCS) assay for detection of unbound copper. Proteins (4 µM) were incubated with $CuSO_4$ (8 µM) in 50 mM sodium phosphate buffer (pH 6.0), with or without ascorbic acid, followed by ultrafiltration (3 kDa MWCO) to remove unbound copper. Copper in the filtrate was quantified using BCS after reduction with ascorbic acid. Fluorescence (Ex/Em 290/325 nm) was converted to concentration using a $CuSO_4$ standard curve. Bound Cu(I) or Cu(II) is expressed stoichiometrically (Cu per µM protein). Data represent mean ± SD (n = 3).

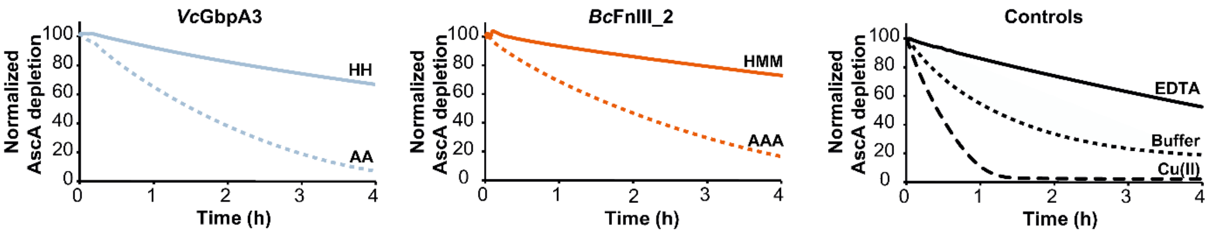


**Figure S11. Probing the ability of GbpA3 and FnIII_2 to bind trace amounts of copper.** Reactions contained 1.33 µM *Vc*GbpA3 or *Bc*FnIII_2 (wildtype and mutant), 1.33 µM EDTA or 1.33 µM Cu(II) as final concentrations in 50 mM sodium phosphate, pH 7.5. To allow binding of Cu(II), mixtures were pre-incubated for 5 min at 37°C. Then, ascorbic acid was added to a final concentration of 1 mM, followed by 10 s of shaking. Absorbance was measuring at 255 nm every 30 s for 4 h. Each reaction was conducted in technical triplicates (n=3); for clarity, only one representative curve per protein is shown, as all replicates were essentially identical.
